# Evolutionary expression patterns in developing biofilms of uropathogenic *Escherichia coli* reveal embryo-like features

**DOI:** 10.1101/2025.04.04.647276

**Authors:** Anja Tušar, Sara Koska, Momir Futo, Niko Kasalo, Ema Svetličić, Nina Čorak, Mirjana Domazet-Lošo, Damjan Franjević, Ivan Mijaković, Tomislav Domazet-Lošo

**Affiliations:** Laboratory of Evolutionary Genetics, Division of Molecular Biology, Ruđer Bošković Institute, Bijenička cesta 54, Zagreb, Croatia; School of Medicine, Catholic University of Croatia, Ilica 244, Zagreb, Croatia; The Novo Nordisk Foundation Center for Biosustainability, Technical University of Denmark, Kgs. Lyngby, Denmark; Faculty of Electrical Engineering and Computing, University of Zagreb, Unska 3, Zagreb, Croatia; Evolution Lab, Department of Biology, Faculty of Science, University of Zagreb, Rooseveltov trg 6, HR-10000 Zagreb, Croatia; Systems and Synthetic Biology Division, Department of Biology and Biological Engineering, Chalmers University of Technology, Gothenburg, Sweden

**Keywords:** biofilms, uropathogenic *Escherichia coli*, development, multicellularity, phylotranscriptomics, phyloproteomics

## Abstract

Developing biofilms of the gram-positive, soil-dwelling bacterium *Bacillus subtilis* exhibit expression patterns and evolutionary imprints similar to those observed in eukaryotic embryos. However, the universality of these evolutionary and developmental regularities has not yet been explored in other biofilm-forming bacteria, including pathogens. To broaden the perspective on ontogeny-phylogeny correlations in bacteria, we recovered phylotranscriptomic and phyloproteomic trajectories throughout the entire biofilm development of uropathogenic *Escherichia coli* UTI89. Here, we show that biofilm growth in *E. coli* UTI89 positively correlates with the evolutionary age of expressed genes, with developing biofilms progressively expressing younger and more divergent genes. These findings suggest that biofilm formation in gram-negative bacteria is not macroevolutionarily naïve and that evolutionary imprints are a pervasive feature of bacterial biofilm development. While these regularities apply to the developmental expression patterns of bulk biofilms, the question remains whether evolutionary stratification occurs within the spatial regions of individual biofilms. To address this, we analyzed expression profiles across concentric regions of *E. coli* UTI89 biofilms at three developmental stages. This analysis revealed that gene expression, functional patterns, and evolutionary imprints are influenced by both developmental timing and distance from the biofilm center. Together, these findings demonstrate that the growth of uropathogenic *E. coli* biofilms is governed by both temporal and spatial macroevolutionary logic, drawing intriguing parallels to organismal development in multicellular eukaryotic lineages.

## Introduction

Multicellular behaviours are an important part of the bacterial life cycle (Shapiro, 1998; Lyons and Kolter, 2015; Flemming and Wuertz, 2019). However, it is not yet clear to what degree bacteria and complex eukaryotes, such as animals, plants, and fungi, share the basic principles of multicellular development (Claessen et al., 2014; Penesyan et al., 2021; Jo et al., 2022). Aggregates of bacterial cells, known as biofilms (Flemming et al., 2021), are particularly relevant for comparison with multicellular eukaryotes, as they represent the most common form of bacterial existence (Flemming and Wuertz, 2019) and exhibit a multitude of collective phenotypes (Claessen et al., 2014; Penesyan et al., 2021; Jo et al., 2022; Sauer et al., 2022).

In our recent study (Futo et al., 2021), we applied phylotranscriptomic and phyloproteomic approaches (Domazet-Lošo and Tautz, 2010b; Futo et al., 2021; Koska et al., 2025) to agar-grown *Bacillus subtilis* biofilms, which are known for their remarkably high structural organization (van Gestel et al., 2015; Ackermann et al., 2015). Our analyses demonstrated that the ontogeny of *B. subtilis* biofilms follows a highly regulated multicellular developmental process, organized into distinct ontogenetic stages that resemble the progression of early developmental stages in animals (Futo et al., 2021).

A striking finding is that *B. subtilis* biofilm development follows a “recapitulation” evolutionary pattern, in which evolutionarily younger and more divergent genes are progressively expressed during the later stages of biofilm growth (Domazet-Lošo and Tautz, 2010b; Futo et al., 2021). This led us to propose that biofilm formation is a bona fide developmental process that, at the systems level, resembles the formation of eukaryotic embryos (Futo et al., 2021). Since a universal strategy of biofilm formation has been suggested across bacteria (O’Toole et al., 2000; Stoodley et al., 2002; Lappin-Scott et al., 2014; Sauer et al., 2022), we hypothesized that the correlation between ontogeny and phylogeny is not unique to gram-positive, non-pathogenic, and soil-dwelling *B. subtilis* biofilms, and that similar evolutionary patterns may also be observed in other biofilm-forming bacteria.

Interestingly, recent studies have revealed that *E. coli* cells clonally organize into rosettes, initiating multicellular development that culminates in the formation of biofilms (Puri et al., 2022; Puri et al., 2024). This discovery is particularly significant, as it suggests that *E. coli* possesses a multicellular life cycle rooted in rosette-initiated development—an adaptation previously unreported outside multicellular eukaryotes (Puri et al., 2022; Puri et al., 2024). This developmental process, guided by rosette formation, seems to be a widespread multicellular phenomenon in *E. coli*, including uropathogenic strains (Puri et al., 2022; Justice et al., 2024).

The majority of uropathogenic *E. coli* isolates can form both extracellular and intracellular biofilms (Whelan et al., 2020; Mukane et al., 2022; Zagaglia et al., 2022). This biofilm-forming capacity is crucial to *E. coli*’s pathogenicity, as it provides protection against host immune responses and promotes the persistence and recurrence of urinary tract infections by employing various biofilm-associated virulence factors (Reigstad et al., 2007; Rijavec et al., 2008; Tajbakhsh et al., 2016; Zagaglia et al., 2022). All of these multicellular features make uropathogenic *E. coli* biofilms particularly suitable for testing whether phylogeny-ontogeny correlations and other system-level analogies to multicellular eukaryotes exist beyond the biofilm development of *B. subtilis*. However, a comprehensive and synchronized global transcriptome and proteome analysis of uropathogenic *E. coli* biofilm development, to our knowledge, has never been performed, making this research effort a necessary first step.

The macroevolutionary imprints identified in *Bacillus subtilis* biofilms were exclusively recovered through the evolutionary analysis of expression profiles across entire developing biofilms (Futo et al., 2021). However, expression patterns can also be studied within distinct spatial regions of developing biofilms. In multicellular eukaryotes, phylotranscriptomic tools are increasingly being applied in the spatial context of developing organisms (Damatac et al., 2024; Ma and Zheng, 2023; Wong et al., 2025). Although spatial heterogeneity plays a critical role in the community organization and function of biofilms (Dragoš et al., 2018; Serra et al., 2013), influencing key processes such as antibiotic resistance and virulence (Silva et al., 2016; Andersson et al., 2019; El-Halfawy et al., 2015), phylotranscriptomic and phyloproteomic analyses have never been applied in this context. While some studies have characterized the spatial organization of transcription within *E. coli* biofilms (Wang et al., 2023; Díaz-Pascual et al., 2021; Beebout et al., 2019), they have focused on a single developmental time point rather than capturing temporal dynamics during biofilm development.

To address this gap, here we recovered spatio-temporal transcriptome and proteome data covering the full developmental process of *E. coli* UTI89 biofilm formation on agar plates. Using this dataset, we investigated ontogeny–phylogeny correlations and functional patterns throughout the development of uropathogenic *E. coli* biofilms. Our findings demonstrate that phylogeny–ontogeny correlations are clearly present in *E. coli* biofilms, reinforcing the idea that macroevolutionary logic is embedded in biofilm development across bacterial species and environmental conditions. Additionally, we show that *E. coli* biofilms exhibit a high degree of spatio-temporal organization, suggesting a striking convergence between biofilm formation and developmental processes in multicellular eukaryotes.

## Results

### *E. coli* biofilms exhibit stage-organized transcription and translation

The ontogeny of multicellular eukaryotes, as well as *B. subtilis* biofilms, is not a continuous process; rather, it is organized into more or less discrete stages, each underpinned by specific expression profiles (Monds and O’Toole, 2009; Levin et al., 2012; Yanai, 2018; Futo et al., 2021). To determine whether *E. coli* UTI89 biofilm growth is also a stage-organized process, we sampled biofilms at 12 time-points covering a full span of biofilm ontogeny, from a liquid starting culture to a 1-month-old biofilms (Figure 1a, Supplementary File S13) and sequenced their transcriptomes.

**Figure 1.**
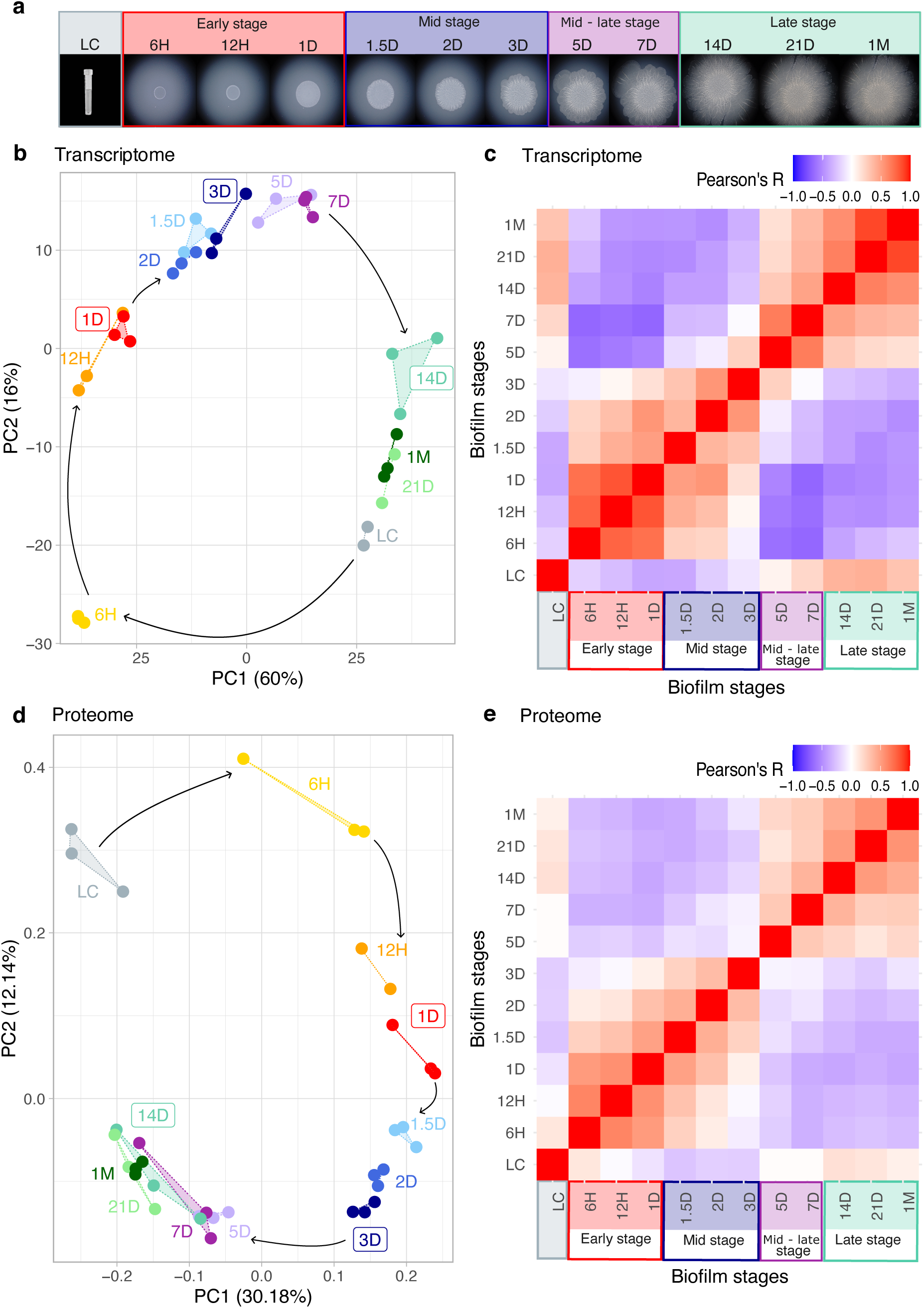
*E. coli* UTI89 biofilm growth exhibits stage-organized gene expression. (**a**) Morphology of *E. coli* UTI89 biofilms on solid agar plates during biofilm maturation at different time points: 6 hours (6H), 12 hours (12H), 1 day (1D), 1.5 days (1.5D), 2 days (2D), 3 days (3D), 5 days (5D), 7 days (7D), 14 days (14D), 21 days (21D), and 1 month (1M) after inoculation with liquid culture (LC). (**b**) Principal component analysis (PCA) of *E. coli* UTI89 transcriptome data and (**d**) proteome data (Supplementary File S1). Biofilm growth time points are represented in different colors. Replicates share the same color and are connected by dotted lines. Black arrows indicate the sequential order of time points in the experimental timeline. Time points framed in boxes (1D, 3D, and 14D) are further analyzed in spatial biofilm experiments. (**c**) Pearson’s correlation coefficients between biofilm ontogeny time points were calculated for transcriptome data, and (**e**) for proteome data (Supplementary File S1). Early (6H–1D), mid (1.5D–3D), mid-late (5D–7D), and late (14D–1M) phases of biofilm growth are depicted in different colors on the x-axis.

In total, we recovered transcriptome data for 4,866 (93%) predicted *E. coli* UTI89 genes, which unveiled a time-resolved principal component analysis (PCA) profile that mirrors the developmental growth trajectory of pathogenic *E. coli* biofilms (Figure 1b). Pearson’s correlation analysis of the transcriptome data further confirmed the presence of distinct developmental stages and identified three main phases of biofilm ontogeny: early (6H–1D), mid (1.5D–3D), and late (14D–1M), with a transitional mid-late stage (5D–7D) (Figure 1c, Supplementary File S1). A similar ontogeny progression was previously observed in the development of non-pathogenic *B. subtilis* biofilms (Futo et al., 2021), supporting the notion that the main strategy of biofilm formation is largely independent of bacterial species (O’Toole et al., 2000; Stoodley et al., 2002; Lappin-Scott et al., 2014; Sauer et al., 2022).

To estimate the overall dynamics of gene regulation, we calculated differential expression across all ontogeny time points, revealing that 4,464 genes (92%) are differentially expressed (Supplementary File S16). When filtering for genes with a minimum twofold expression change, this number decreases to 1,639 (34%). To further explore expression dynamics between successive ontogeny time points, we conducted pairwise transcriptome comparisons (Supplementary Figure 11ab, Supplementary File S17). On average, 7% of expressed genes were significantly upregulated by more than twofold during transitions between successive stages (Supplementary Figure 11ab, Supplementary File S17). Although these expression changes are substantial, they remain lower than those observed during *B. subtilis* biofilm development (Futo et al., 2021).

Similar to *B. subtilis* biofilms (Futo et al., 2021), we detected the most dramatic transcriptional shift upon inoculation of agar plates with liquid culture. This transcriptional change, also visible in the PCA (Figure 1b), occurred during the transition from liquid culture (LC) to the early biofilm lifestyle (6H), where 26% of expressed were significantly upregulated by more than twofold (Supplementary Figure 11ab, Supplementary File S17). These results further confirm that bacterial cells generally undergo substantial transcriptome changes as they transition from a planktonic state to a surface-attached community (O’Toole et al., 2000; Futo et al., 2021; Sauer et al., 2022). The next most dynamic transition in gene upregulation occurred during the late phase of biofilm growth (7D–14D), where approximately 18% of expressed *E. coli* genes were significantly upregulated by more than two fold (Supplementary Figure 11b, Supplementary File S17). Taken together, transcriptome analyses reveal that biofilm development in *E. coli* UTI89 is a non-continuous process, consisting of distinct ontogeny phases (early, mid, mid-late, and late), each characterized by unique expression shifts.

Correlation between transcript and protein levels is generally low (Liu et al., 2016; Futo et al., 2021), highlighting the need to assess both molecular phenotypes to obtain a comprehensive picture of gene expression. However, state-of-the-art protein quantification via mass spectrometry has lower genome-wide coverage and reduced sensitivity for low-abundance proteins compared to transcriptome sequencing (Buccitelli and Selbach, 2020). While transcriptome sequencing provides a more reliable representation of gene expression dynamics, integrating both techniques offer a more complete view, linking transcriptional regulation with protein-level function (Futo et al., 2021; Buccitelli and Selbach, 2020).

We thus quantified the proteomes of the same *E. coli* UTI89 biofilm stages that we sampled for transcriptome sequencing (Figure 1a). Our proteomic analysis identified 2,518 predicted proteins, covering 48% of the theoretical *E. coli* UTI89 proteome. Although this represents nearly half the number of genes recovered by transcriptome sequencing, principal component analysis (PCA) and Pearson’s correlation analyses revealed largely congruent patterns between the transcriptome and proteome (Figure 1d,e), with a time-resolved succession of biofilm stages (Figure 1d) and distinct ontogeny phases (Figure 1e). These findings support the ontogeny progression detected in the transcriptomic data (Figure 1b,c), demonstrating that biofilm development follows a structured, stage-specific pattern at both the transcriptomic and proteomic levels.

Compared to transcriptome analysis, proteome analysis exhibited fewer differentially expressed proteins across all time points of biofilm ontogeny (1,301 proteins, 68%) (Supplementary File S16). However, the proportion of upregulated proteins displaying at least a two-fold expression change (711 proteins, 37%) was comparable to that observed in the transcriptome dataset. In pairwise comparisons between biofilm stages an average of 8% of proteins exhibited upregulated differential expression with a minimum twofold change (Supplementary Figure 11b, Supplementary File S8), a lower percentage than observed at the transcriptomic level (Supplementary Figure 11a). Consistent with prior transcriptomic findings (Supplementary Figure 11ab, Supplementary File S17) the most pronounced transcriptional shift occurred during the transition from liquid culture (LC) to the early biofilm stage (6H) (Supplementary Figure 11cd, Supplementary File S8). Collectively, the proteomic data also delineate distinct ontogeny phases and the structured progression of biofilm ontogeny, despite the generally low correlation between the transcriptome and proteome data and the inherent limits of protein quantification (Supplementary Figure 5).

### Ontogeny-phylogeny correlations in *E. coli* biofilm development

To explore whether *E. coli* UTI89 biofilm development follows an evolutionary logic in the form of phylogeny–ontogeny correlations—similar to *B. subtilis* biofilms and multicellular eukaryotes (Futo et al., 2021; Domazet-Lošo and Tautz, 2010a; Quint et al., 2012)—we linked transcriptome and proteome profiles to evolutionary gene age estimates, which we obtained using a phylostratigraphic approach (Domazet-Lošo et al., 2007; Futo et al., 2021; Čorak et al., 2023). To obtain gene age estimates, we first constructed a consensus phylogenetic tree that begins at the origin of cellular organisms and ends with *E. coli* UTI89 as the focal strain. We then mapped 5,206 (99.9%) *E. coli* UTI89 genes to nine phylostrata (ps), with ps1 representing the evolutionary oldest and ps9 representing the evolutionary youngest phylostratum (Supplementary Figure 1, Supplementary File S21, Supplementary File S22).

Using these evolutionary gene age estimates along with transcriptome and proteome expression values, we calculated the transcriptome age index (TAI) and the proteome age index (PAI) (Figure 2, Supplementary File S4). TAI and PAI are cumulative measures that reflect the overall evolutionary age of the expressed genes at a particular stage of biofilm development (Domazet-Lošo and Tautz, 2010a; Futo et al., 2021). Similar to *B. subtilis* biofilms (Futo et al., 2021), *E. coli* UTI89 biofilms exhibited a positive correlation between biofilm developmental timing and the evolutionary age of expressed genes at both the transcriptome (Figure 2a) and proteome (Figure 2b) levels. In both analyses, evolutionarily younger genes are increasingly expressed as the biofilm progresses to later developmental stages (Figure 2a,b). This type of regularity is known as the ‘recapitulation’ pattern, originally proposed in the 19th century as a general feature of animal development. In contemporary studies, it is often referred to as the early conservation model (Futo et al., 2021). Interestingly, in late-stage biofilms at the transcriptome level an opposite trend emerges, with transcriptomes becoming progressively older between 7D and 1M stages (Figure 2a). This is another analogy to animals and plants which also show a drop in TAI during the aging process (Domazet-Lošo and Tautz, 2010b, Koska et al. 2025).

**Figure 2.**
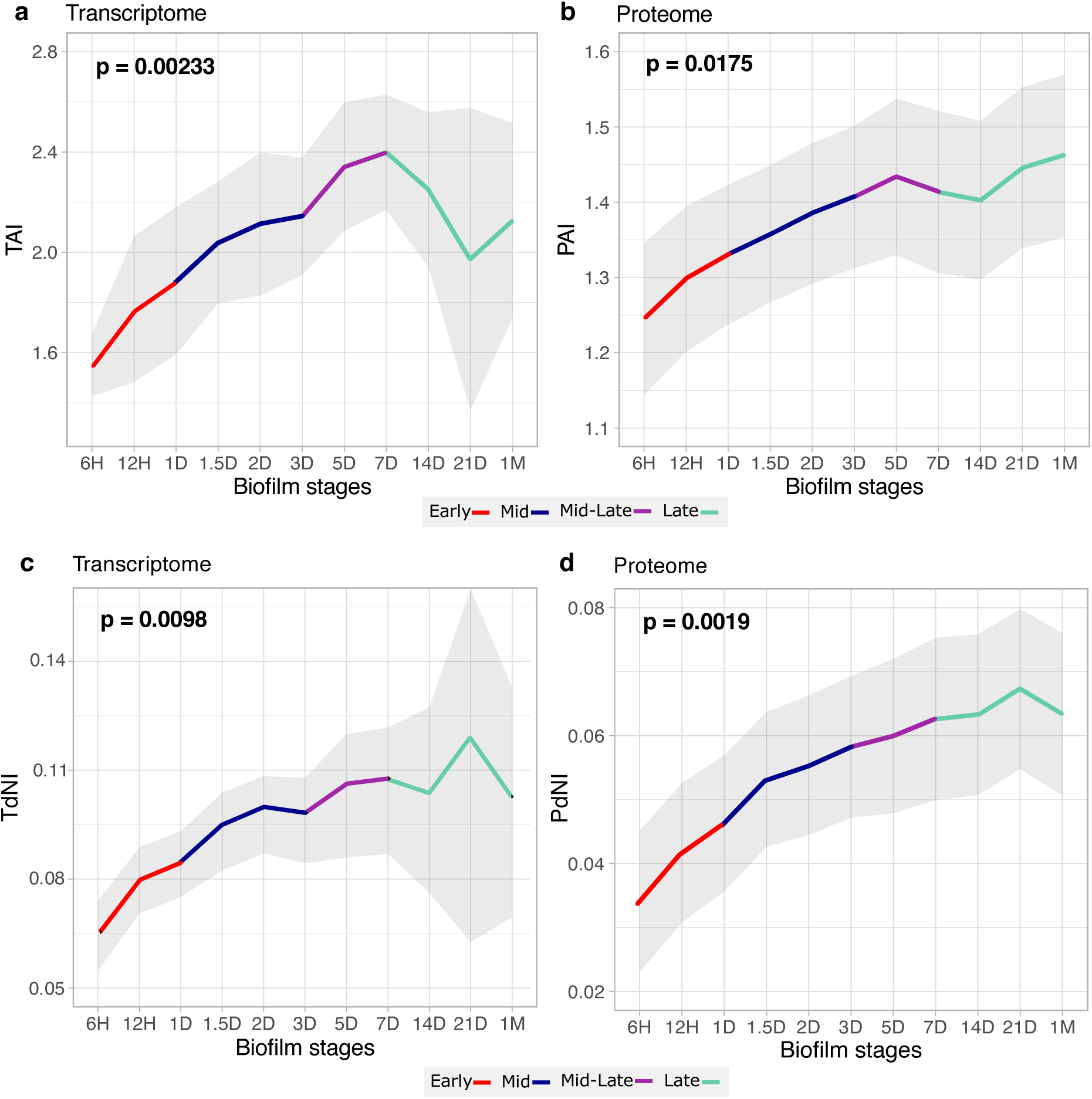
Ontogeny recapitulates phylogeny in uropathogenic *E. coli* biofilms. (**a**) Transcriptome age index (TAI) and (**b**) proteome age index (PAI) profiles of *E. coli* UTI89 biofilm ontogeny reveal a recapitulation pattern, where early biofilm stages predominantly express evolutionarily older genes, while later stages express younger genes (Supplementary File S4). (**c**) Transcriptome non-synonymous divergence index (TdNI) and (**d**) proteome non-synonymous divergence index (PdNI) also reveal a recapitulation pattern, where early biofilm stages predominantly express evolutionarily conserved genes, while later stages express more diverged genes at non-synonymous sites (Supplementary File S6). Non-synonymous divergence rates were estimated by comparing *E. coli* UTI89 coding sequences to *Klebsiella pneumoniae* (Enterobacteriaceae). Pairwise comparison to more divergent *Yersinia pestis* (Enterobacterales), showed similar results (Supplementary Figure 2, Supplementary File S7). We evaluated significance (p-value) of these evolutionary patterns using early conservation test (see Methods). Grey shaded areas indicate ± one standard deviation based on permutation analysis. Different biofilm developmental stages are color-coded: early (red), mid (blue), mid-late (purple), and late (green).

TAI and PAI measures, which are based on the phylostratigraphic approach, estimate the cumulative evolutionary age of transcriptomes and proteomes by identifying relatively remote homologs across the tree of life (Domazet-Lošo et al., 2007; Domazet-Lošo and Tautz, 2010a; Futo et al., 2021; Domazet-Lošo et al., 2024). However, an alternative approach to analyzing evolutionary imprints in expression datasets involves estimating the evolutionary divergence rates of coding sequences, rather than the age of entire genes (Quint et al., 2012; Futo et al., 2021; Koska et al., 2025). In this context, to account for codon usage bias, we recently introduced the transcriptome and proteome nonsynonymous (TdNI, PdNI) and synonymous (TdSI, PdSI) divergence indices, which rely on nonsynonymous and synonymous divergence rates, respectively (Futo et al., 2021).

To estimate nonsynonymous (dN) and synonymous (dS) divergence rates, we aligned the coding sequences of *E. coli* UTI89 with those of the relatively closely related *Klebsiella pneumoniae* MGH78578. When analyzing the transcriptome dataset, both TdNI (Figure 2c) and TdSI (Supplementary Figure 3) exhibited a significant recapitulation pattern, with the divergence rates of transcribed genes—at both nonsynonymous and synonymous sites— progressively increasing toward the later stages of biofilm growth. A comparable pattern was observed when examining nonsynonymous (PdNI) and synonymous (PdSI) divergence rates in the proteomic context (Figure 2c, Supplementary Figure 3). To test the robustness of these results regarding the choice of bacterial species, we repeated the analyses using more divergent *Yersinia pestis,* within the gram-negative order Enterobacterales, which yielded highly congruent results (Supplementary Figure 3).

In line with the patterns observed for synonymous divergence rates, codon usage bias— assessed by the transcriptome codon usage index (TCBI) and proteome codon usage index (PCBI)—also exhibited a significant recapitulation pattern, with progressively lower bias toward the later stages of biofilm development (Supplementary Figure 3). Together, transcriptome and proteome evolutionary indices suggest that similar evolutionary constraints shape developing biofilm transcriptomes at both deep (TAI, PAI) and more recent (TdNI, TdSI, TCBI, PdNI, PdSI, PCBI) evolutionary levels. This results in a recapitulation pattern, where more divergent genes are progressively expressed at higher levels in the later stages of biofilm growth.

### *E. coli* biofilm development follows a functionally punctuated architecture

To analyze functional patterns during *E. coli* UTI89 biofilm development, we performed a functional enrichment analysis of transcripts and proteins using GO annotations (Figure 3, Supplementary Figure 6). The enrichment analysis of GO annotations revealed tight functional control over biofilm development, where each biofilm stage expresses a distinct set of functions (Figure 3, Supplementary file 7). This pattern parallels *B. subtilis* biofilm development, which also displays a progression of functionally discrete stages (Futo et al., 2021). Interestingly, genes with unknown functions are significantly enriched in transcriptomes from mid to late stages of biofilm development (2D to 1M), suggesting that key molecular mechanisms underlying *E. coli* UTI89 biofilm development remain incompletely understood (Figure 3). Statistical analysis of unannotated genes on the phylostratigraphic map revealed that they are increasingly enriched from Bacteria (ps2) to *E. coli* (ps9) as focal species, indicating that they are more common in younger phylostrata (Supplementary Figure 4). This resembles development in animals where phylogenetically restricted genes (orphans) are expressed in later stages of ontogeny where they contribute to the generation of clade specific phenotypes (Domazet-Lošo and Tautz, 2010a; Tautz and Domazet-Lošo, 2011; Xia et al. 2025).

**Figure 3.**
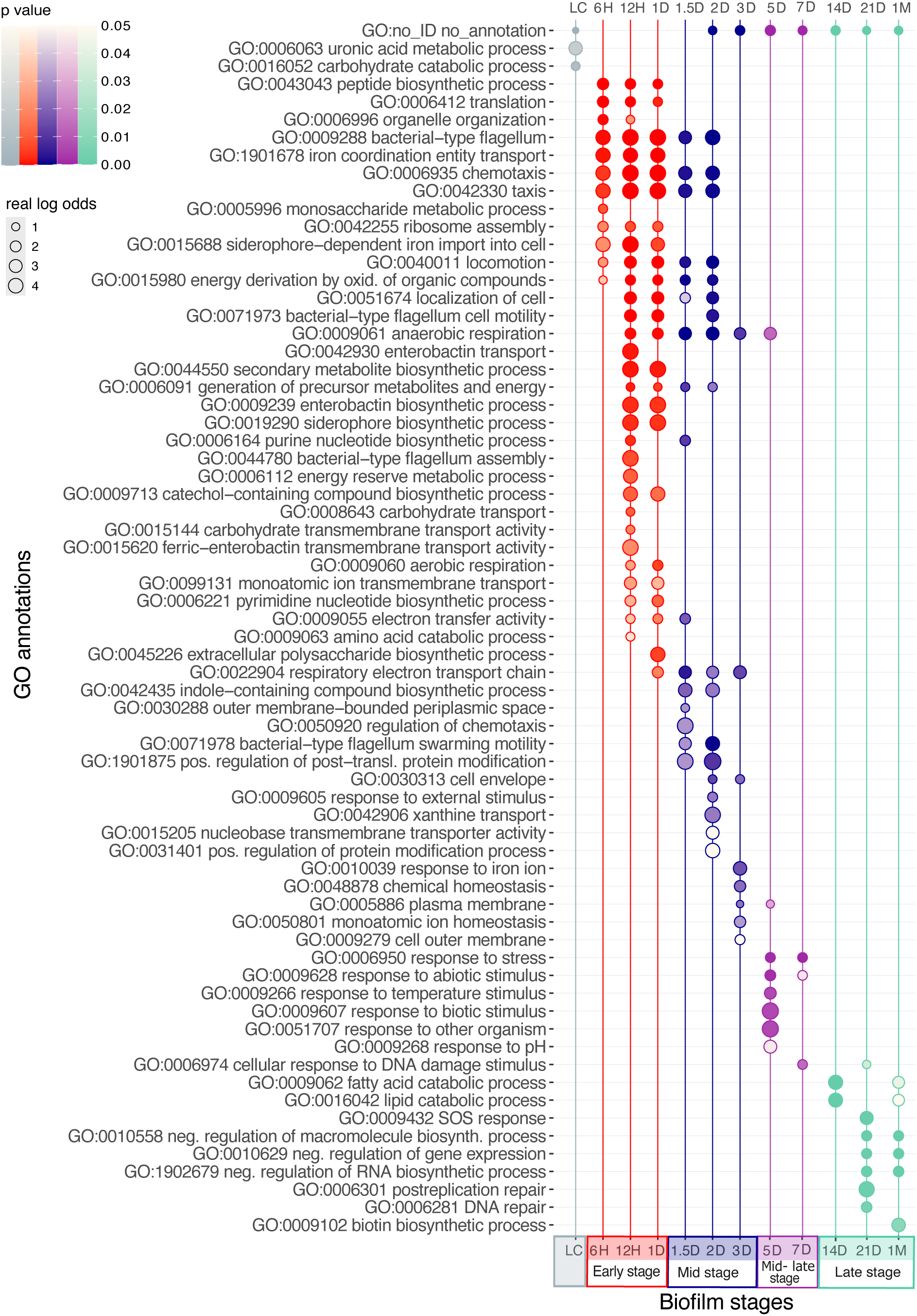
*E. coli* UTI89 biofilm development is organized into functionally discrete stages. Functional enrichment analysis of GO annotations was performed for genes expressed at least 0.5 times (log2 scale) above the median of their overall transcription profile at a given stage (Supplementary File S14). The significance of enrichment was assessed using a one-tailed hypergeometric test, corrected for multiple comparisons using the Benjamini–Hochberg procedure. Only functional enrichments with a p-value below 0.05 were considered. The complete set of enriched GO annotations is listed in Supplementary File S3, while this figure displays only a non-redundant set of GO functions. The magnitude of functional enrichments is represented by log odds (circle sizes), while p-value significance is indicated by color shading. The stages of biofilm development, which include the starting culture (grey), early phase (red), mid phase (blue), mid-late phase (purple), and late phase (green), are color-coded.

During the early stages of biofilm growth (6H–1D), the enriched functions were primarily associated with biosynthetic processes, energy generation, ribosomal synthesis, iron homeostasis, and cell motility—all of which are indicative of active biofilm formation (Figure 3). In the mid-phase (1.5D–3D), functions related to locomotion, bacterial-type flagellum, and energy production were enriched, suggesting active biofilm spreading on agar plates (Figure 3). In the mid-late (5D–7D) and late-phase (14D–1M), enriched functions shifted toward the regulation of metabolic processes, post-replication repair, and stress response (Figure 3). For instance, the enrichment of functions related to the SOS response and DNA repair at 21D suggests harsh environmental conditions that biofilms face at this stage (Figure 3). Proteome-level functional enrichment analyses exhibit stage-specific enrichment patterns similar to those observed at the transcriptome level (Supplementary Figure 6). This demonstrates that at the transcriptome as well as proteome level, despite their poor correlation (Supplementary Figure 5), *E. coli* UTI89 biofilm growth displays a stepwise functional architecture.

Recently discovered multicellular life cycle in *E. coli* that is initialed by developmental rosettes is also underpinned by a tight transcriptional regulation (Puri et al., 2022; Puri et al., 2024). It is therefore interesting to look for expression patterns of genes that are implicated in the rosette-initiated life cycle (Puri et al., 2022; Puri et al., 2024). The expression profiles for these genes display temporal succession that roughly correspond to developmental regulation originally described for rosette-initiated life cycle (Supplementary Figure 12, Supplementary Figure 13). For example, genes related to type-1 fimbriae and flagella peak in expression in early biofilms, while genes related to curli peak in mid-stage biofilms (Supplementary Figure 12, Supplementary Figure 13).

Taken together, functional patterns indicate that *E. coli* biofilm development follows a discrete organization with fine temporal grading that aligns with transcriptome and proteome stages. Similar to animals (Domazet-Lošo and Tautz, 2010a), plants (Quint et al., 2012; Koska et al., 2025), and *B. subtilis* (Futo et al., 2021), *E. coli* biofilm development exhibits a distinct, and stage-organized developmental architecture (Levin et al., 2012; Yanai, 2018; Futo et al., 2021).

### Spatio-temporal transcriptomes and proteomes of *E. coli* UTI89 biofilm development

To gain insight into the global spatiotemporal regulation of gene transcription and protein translation during *E. coli* biofilm development, we sequenced transcriptomes and quantified proteomes from different biofilm regions at three time points (Figure 4a). Specifically, we collected inner and outer biofilm regions from 1D (early-phase) and 3D (mid-phase) biofilms, as well as inner, middle, and outer regions from 14D (late-phase) biofilms. In total, spatial samples yielded transcriptome data for 4,989 (∼96%) and proteome data for 2,355 (45.2%) *E. coli* UTI89 genes (Supplementary File S15). Principal component analysis (PCA) revealed distinct transcriptomic and proteomic profiles across biofilm regions, with biological replicates clustering closely together (Figure 4b,d). However, along with Pearson’s correlation coefficients from an all-against-all comparison (Figure 4c,e), PCA indicated that the temporal progression of biofilm growth is a stronger determinant of gene expression than spatial positioning within biofilms.

**Figure 4.**
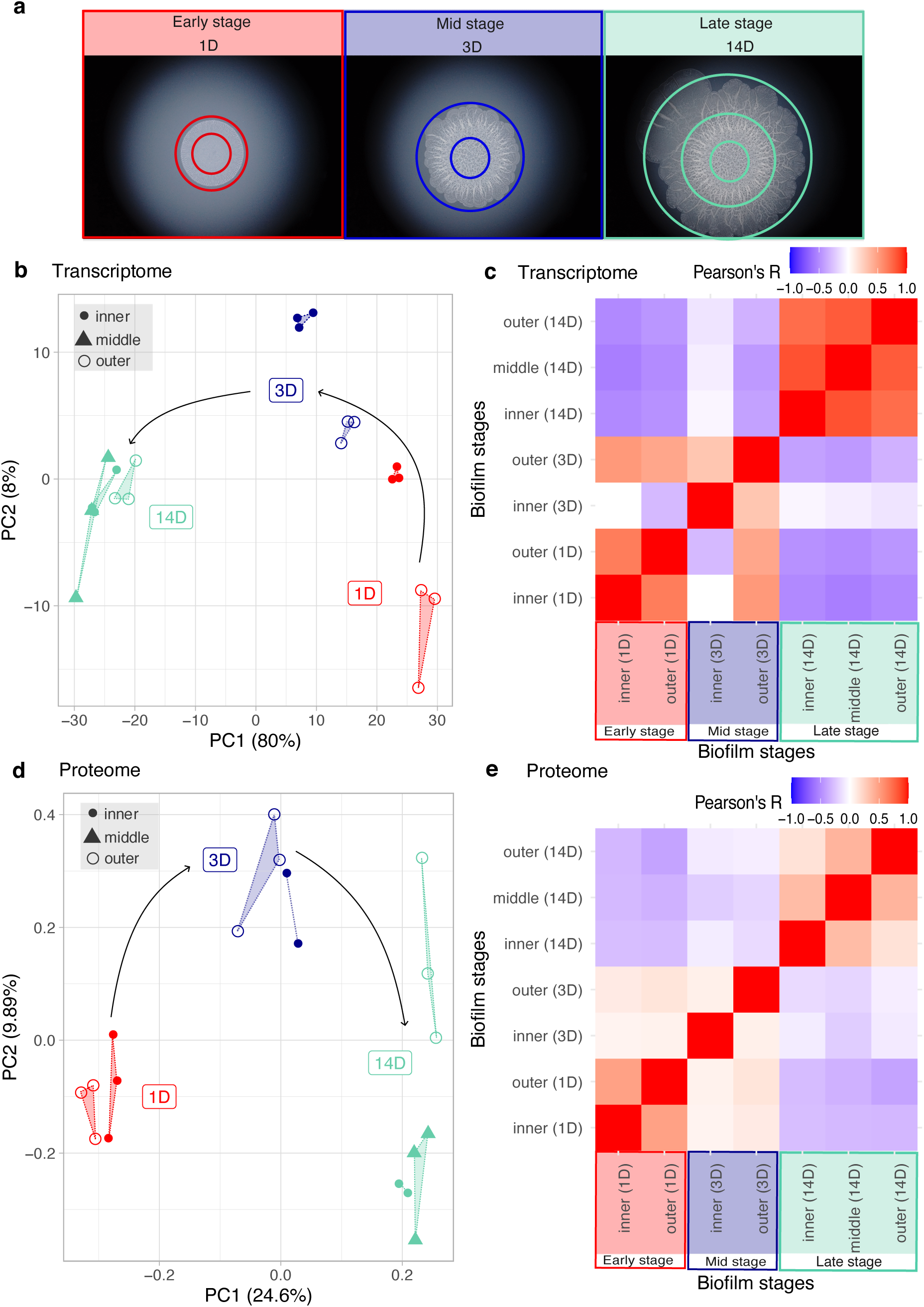
*E. coli* UTI89 biofilms exhibit spatially and temporally organized gene expression. (**a**) Schematic representation of sampling regions in 1D (early-stage), 3D (mid-stage), and 14D (late-stage) biofilms. Sampling locations—inner, middle, and outer biofilm regions—were determined based on biofilm morphology and are depicted by circles. (**b, d**) Principal component analysis (PCA) of the recovered (**b**) transcriptome and (**d**) proteome data (Supplementary File S15). Biofilm stages are color-coded: 1D (red), 3D (blue), and 14D (green). Biofilm regions are represented by symbols: inner region (filled circle), middle region (filled triangle), and outer region (empty circle). Biological replicates are connected by dotted lines, and black arrows indicate the chronological progression of biofilm stages in the experimental timeline. (**c, e**) Pearson’s correlation coefficients from an all-against-all comparison of (**c**) transcriptome and (**e**) proteome data across biofilm regions and developmental time points (Supplementary File S15). The x-axis color-codes early (1D), mid (3D), and late (14D) stages for clarity.

Among genes that were expressed in spatial samples 32% (1D), 44% (3D), and 9% (14D) showed significant change of expression between spatial regions (Supplementary File S24), indicating considerable spatial regulation within the biofilm. Interestingly, the greatest differentiation between inner and outer regions was observed in 3D biofilms, where the outer region of 3D biofilms were more similar to both inner and outer regions of 1D biofilms than to the inner region of 3D biofilms (Figure 4c,e). To further explore this spatial heterogeneity of expression patterns in biofilms, we performed functional analyses of differentially transcribed genes, revealing spatially organized functional distribution in *E. coli* UTI89 biofilms (Supplementary Figures 7–10).

The outer region of the 1D biofilms is primarily enriched in transcripts related to iron acquisition systems, ribosome complex synthesis, translation, cell motility, and carbohydrate metabolism. In contrast, the inner region is predominantly associated with amino acid biosynthesis and stress responses (Supplementary Figure 7). Similarly, in 3D biofilms, the outer region is enriched in functions related to nucleotide production, amino acid catabolism, and transporter systems. On the other hand, the inner region of mid-stage (3D) biofilms, which is transcriptionally the most distinct (Figure 4c), is enriched with genes of unknown function and those responsible for social and environmental interactions (Supplementary Figure 8).

In late-stage (14D) biofilms, the outer region is enriched in functions related to polysaccharide metabolism, biofilm formation, cellular pH regulation, and protein folding. Compared to the outer region, the middle region exhibits increased transporter activity, while the inner region is primarily associated with amino acid catabolism (Supplementary Figure 9). At the proteome level, we observed functional patterns similar to those at the transcriptome level. However, the magnitude of significant changes in protein expression between spatial regions within the biofilm was much lower than at the transcriptome level—6% at 1D, 7% at 3D, and 10% at 14D (Supplementary File S26)—likely due to the lower coverage and dynamic range of the protein dataset (Supplementary Figure 10).

These findings suggest that the outer and inner regions of biofilms play distinct functional roles in biofilm development. Moreover, some metabolic functions are not strictly confined to a specific spatial region but can shift between regions depending on the stage of biofilm development. For example, amino acid production is localized to the inner region of early-stage (1D) biofilms, while amino acid catabolism predominates in the outer region of mid-stage (3D) biofilms. Notably, the functions related to amino acid catabolism shift to the inner region in late-stage (14D) biofilms.

### Evolutionary imprints in biofilm regions

Given that genes with unknown function have a more recent evolutionary origin (Supplementary Figure 4) and that genes with unknown function are also enriched in the inner part of the mid-stage 3D biofilms (Supplementary Figure 8), we suspected that spatial regions of *E. coli* UTI89 biofilms will also differ between each other in the terms of macroevolutionary imprints. To investigate this, we compared TAI and PAI values across different biofilm regions (Figure 5). In 1D (early-stage) biofilms, we observed a non-significant trend where the inner region expressed a comparatively younger transcriptome and proteome (Figure 5a, d). Nevertheless, this trend became stronger and statistically significant in more mature 3D (mid-stage) biofilms (Figure 5b, e). Interestingly, in 14D (late-stage) biofilms, this trend reversed in transcriptomes, with the inner region exhibiting an evolutionarily older transcriptome, while the outer region displayed a younger transcriptome (Figure 5c). However, this transcriptomic shift was not reflected in proteomes (Figure 5f), which is predictable given the lower coverage and dynamic range of proteome data. Together, these findings reveal that macroevolutionary patterns are present not only along the temporal trajectory of biofilm growth but also between inner and outer biofilm regions, particularly in well-developed, mid-stage biofilms.

**Figure 5.**
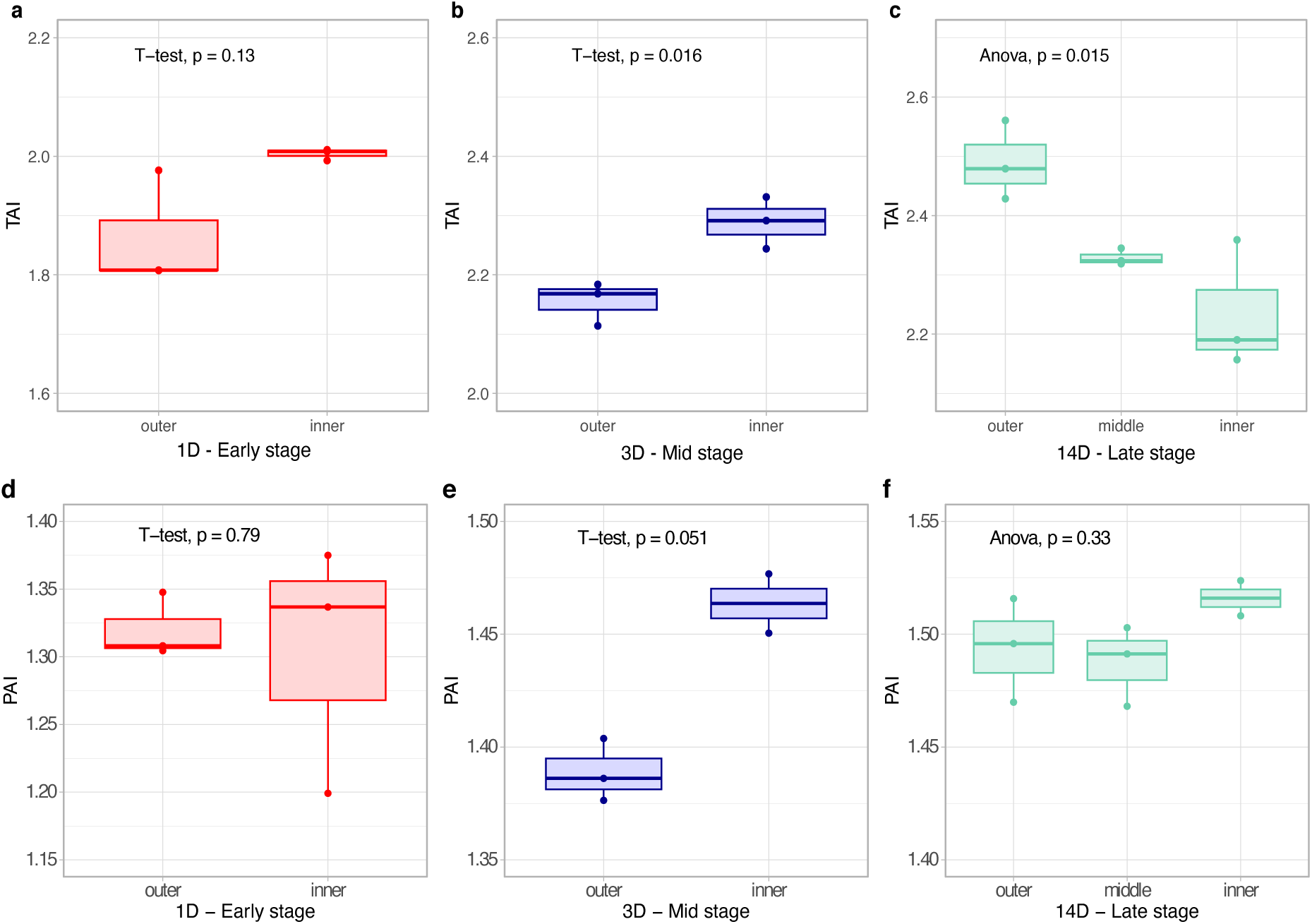
Comparison of evolutionary indices between inner and outer *E. coli* UTI89 biofilm regions. (**a–c**) Transcriptome age index (TAI) comparison between inner and outer biofilm regions in (**a**) 1D, (**b**) 3D, and (**c**) 14D biofilms (Supplementary File S18). (d–f) Proteome age index (PAI) comparison between (**d**) 1D, (**e**) 3D, and (**f**) 14D biofilms (Supplementary File S18). The significance of TAI and PAI differences between biofilm regions was assessed using a t-test for (**a, b**) and ANOVA for (**c**). Statistical tests were performed with three biological replicates per sample. Replicates are depicted as individual dots in the boxplots.

## DISCUSSION

This study revealed that the developmental dynamics of gene expression in uropathogenic *E. coli* UTI89 biofilms parallel analogous developmental processes in animals (Domazet-Lošo and Tautz, 2010), plants (Koska et al., 2025), and *B. subtilis* (Futo et al., 2021). Despite their independent origins and vast evolutionary distances on the tree of life, these diverse developmental systems nevertheless exhibit striking system-level analogies, including punctuated expression dynamics, spatiotemporal functional stratification, and ontogeny-phylogeny correlations. Recent research suggests that these analogies extend as far as the amino acid anabolism and the evolution of behavioral patterns, as exemplified by convergent events in animals and bacteria (Kasalo et al. 2024; 2025a; 2025b). Similar to embryonic development, we found that *E. coli* UTI89 biofilms display a structured progression of gene and protein expression across both time and space. These coordinated expression patterns outline functional and evolutionary architecture *E. coli* UTI89 biofilms.

The observation that gene functions shift across biofilm stages supports the notion of biofilm formation as a coordinated, stage-specific multicellular developmental process (Figure 3) (Futo et al., 2021; Puri et al., 2023; Puri et al., 2024). For example, the early phase of *E. coli* UTI89 biofilm growth are characterized by protein synthesis and motility, while in the mid phase function like response to environmental stimuli, communication at interfaces and uncharacterized functions/proteins dominate. On the other hand, the response to stress is a prevailing function activity of the late phase biofilms (Figure 3). This functional sequence in developing *E. coli* UTI89 biofilms roughly resembles the progression of functions in *B. subtilis* biofilm formation (Futo et al., 2021), suggesting that the bacterial life in the form of biofilms is shaped, at independent instances, by similar selective forces (O’Toole et al., 2000; Stoodley et al., 2002; Lappin-Scott et al., 2014; Sauer et al., 2022, Futo et al., 2021). By covering all temporal stages, from initial attachment to the starvation response, this study offers an exhaustive view of *E. coli* UTI89 biofilm development and provides a benchmark for future investigations into biofilm physiology and pathogenesis.

Intriguingly, *E. coli* UTI89 biofilms also show an evolutionary architecture, where developing biofilms progressively express younger and more divergent genes (Figure 2). This macroevolutionary imprint, present also in *B. subtilis* biofilm formation (Futo et al., 2021) and known as a recapitulation pattern, suggests that bacterial biofilms, similar to eukaryotic embryos (Domazet-Lošo and Tautz, 2010, Quint et al., 2012, Koska et al. 2025), harbour a memory on evolutionary past. The parallels between the biofilms of gram-negative *E. coli* and gram-positive *B. subtilis*, both in development and in its evolutionary imprints (Futo et al., 2021), suggest that universal patterns may exist among diverse biofilm-forming bacteria. The existence of the universal biofilm model (Stoodley et al., 2002; Hall-Stoodley et al., 2004) leads us to believe that the ontogeny-phylogeny correlations identified in the biofilm development of *E. coli* and *B. subtilis* are likely to be present in other biofilms, both in situ and ex situ.

Phylostratigraphy tools (TAI, PAI) have been successfully used to detect phylogeny-ontogeny correlations in metazoans (Domazet-Lošo and Tautz, 2010a; Kalinka et al. 2010), plants (Quint et al. 2012, Koska et al. 2025), fungi (Cheng et al. 2015), brown algae (Lotharukpong et al. 2024), *B. subtilis* (Futo et al., 2021) and in this study *E. coli*. Notably, the recapitulation pattern originally proposed for animals is now observed in bacteria (Figure 2). Our results using phylostratigraphy-based tools (TAI, PAI) are fully supported with findings from methodologically independent divergence-based tools (TdNI, TdSI, PdNI, PdSI) (Figure 2, Supplementary Figure 2, Supplementary Figure 3). These positive phylogeny-ontogeny correlations and stage-organized gene expression architecture suggest that biofilm growth in both gram-negative pathogenic *E. coli* and gram-positive non-pathogenic *B. subtilis* represents a true multicellular developmental process (Futo et al., 2021).

In addition, in this study we explored expression patterns in the spatial regions of developing biofilms (Figure 4). In contrast to previous studies which consider only one temporal point of biofilm growth (Wang et al., 2023; Diaz-Pascual, 2021; Beebout et al., 2019), we here analyzed temporal dynamics of spatial regions by covering biofilms in early, mid and late phase of their growth. This allowed us to catch previously unrecognized temporal dynamic within spatial regions, where cells in different regions shift their functional and developmental roles. For instance, the presence of morphologically distinct subpopulations within biofilm communities and the extensive research on spatial organization within biofilms suggest that the expression of biofilm-related genes must be spatially localized (Beebout et al., 2019; Heacock-Kang et al., 2017; Yannarell et al., 2023). In this study, we identified a curious phenomenon where younger genes, involved in biofilm formation, are predominantly expressed in the inner regions of early and mid-stage biofilms, while this trend reverses in late-stage biofilms (Figure 5).

Another unexpected phenomenon regarding the localization of functions in the biofilm is visible in comparison with the study of the non-pathogenic *E. coli* biofilms, where the authors identify heightened expression of genes related to arginine biosynthesis in the periphery of the biofilm, while the arginine utilization genes are primarily expressed in the interior (Wang et al. 2023). We, however, recovered exactly the opposite trend in the early and mid-stages of the biofilm growth (Figure 6). In the late stage, there is no clear spatial segregation in the expression of these genes, indicating that arginine metabolism became more spatially diffused (Figure 6). Since our enrichment results for other functions within inner and outer spatial regions are largely congruent with those in Wang et al. (2023), it is likely that factors such as differences in growth conditions and strain idiosyncrasy determine this disparity.

**Figure 6.**
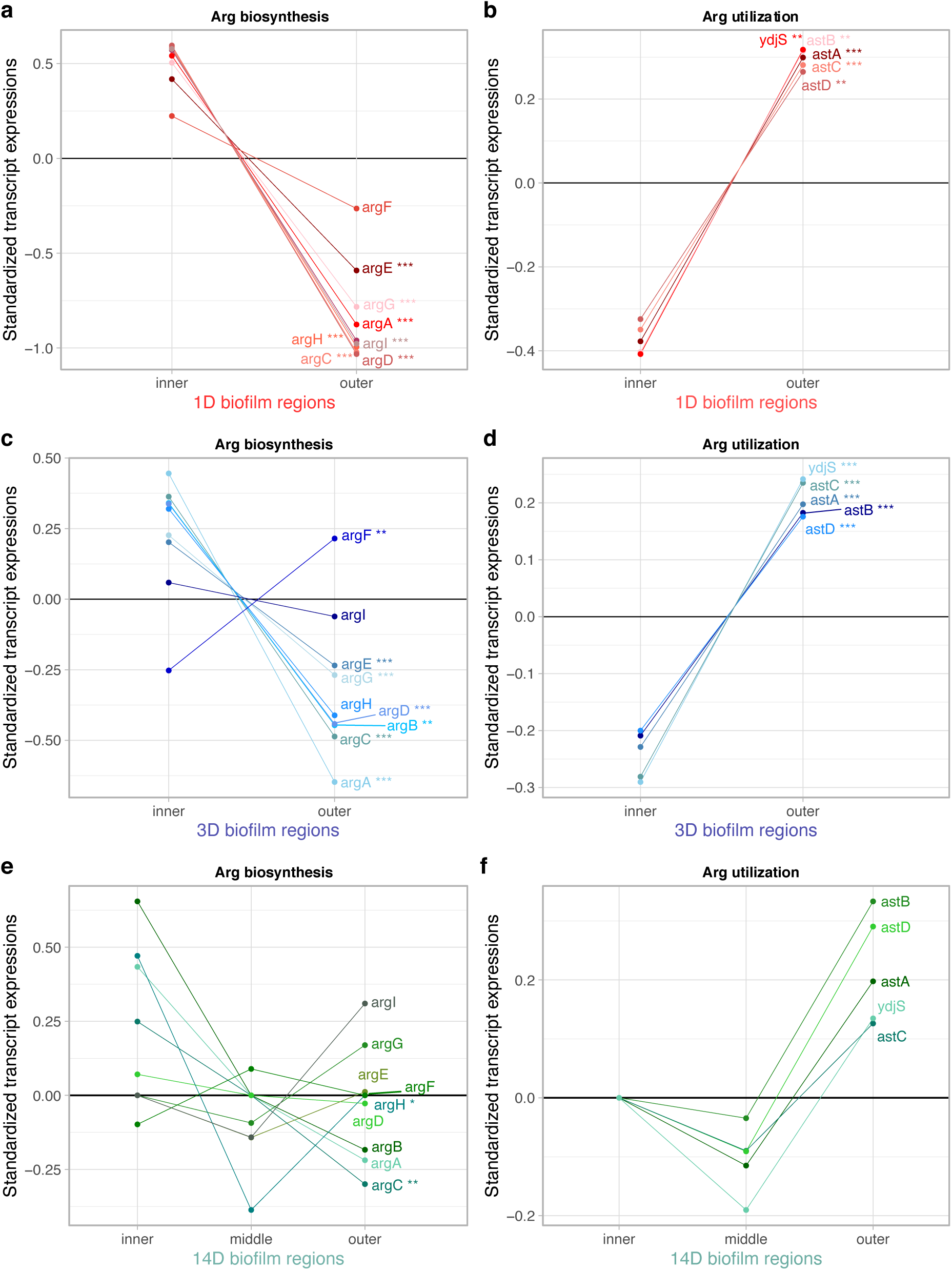
Standardized expression profiles for arginine biosynthesis and utilization genes in the *E. coli* UTI89 biofilm across spatial regions and temporal stages. **(a,b)** early stage (1D), **(c,d)** mid stage (3D), **(e,f)** late stage (14D), **(a,c,e)** arginine biosynthesis genes, **(b,d,f)** arginine utilization genes. To standardize gene expression profiles to the same scale, each gene underwent normalization to the median and log2 transformation, resulting in per-gene standardized expression values (Supplementary file S28). We utilized the DESeq2 V1.38.3 (Love et al., 2014) R package’s pipeline to estimate the overall differential expression of each gene across all biofilm developmental stages and spatial regions within the biofilm, employing the likelihood ratio test (LRT) (Supplementary File S24). Statistically significant results are indicated with asterisks (* p < 0.05, ** p < 0.01, *** p < 0.001).

Although biofilms are increasingly regarded as multicellular organisms (Futo et al., 2021; Puri et al., 2023; Puri et al., 2024; Sauer et al., 2022), the bacterial differentiation within biofilm subpopulations is more flexible and less terminal than the differentiation of embryonic stem cells in animals (Dodson and Kennedy., 2020). This flexibility allows bacteria to rapidly respond to environmental changes, resulting in dynamic subpopulations with reduced reliance on strict spatial differentiation. This could explain the difference in the localization of arginine metabolism between our results and those of Wang et al. (2023). It seems that even the basic metabolic functions, such as amino acid biosynthesis and utilization can shift between spatial regions depending on environmental conditions, e.g., nutrient availability. Another possibility is that pathogenic biofilms have a fundamentally different localization of metabolic functions in comparison to non-pathogenic ones.

These observations underscore the unique adaptive strategies of bacterial biofilms and their distinction from eukaryotic multicellular organisms. Unlike animals, which exhibit precise spatial localization of developmentally relevant gene expression (Tomancak et al., 2007), *E. coli* biofilms show a lesser degree of spatial organization (Figure 4). While distinct transcriptomic and proteomic dynamics exist across biofilm subpopulations, our findings suggest that temporal organization during biofilm ontogeny has a greater impact than spatial differentiation (Figure 1, Figure 4).

In this study, we conducted research under *in vitro* conditions growing *E. coli* biofilms on a solid – air interface, which does not fully replicate the complexity of *in vivo* effects. However, despite this limitation, the broad spectrum of expressed functions captured in our analysis, from fundamental bacterial metabolic pathways to biofilm formation and virulence-related genes, suggests that the observed ontogeny-phylogeny correlations are likely relevant to *in vivo* biofilms. The presence of such correlations in natural settings remains to be explored. It is likely that symbiotic interactions in multi-species biofilms and specific temporal ecological fluctuations introduce additional variations to the general evolutionary trend. Our approach may aid in identifying early regulatory shifts in biofilm development by tracking variations in ontogeny-phylogeny correlations. Additionally, categorizing genes according to their evolutionary origins and roles in biofilm development establishes a framework for uncovering key genes essential to this process. The effectiveness of phylostratigraphy has already been demonstrated in the discovery of novel sporulation genes in *B. subtilis* (Shi et al., 2020). Exploring these options will deepen our understanding of uropathogenic *E. coli* biofilms and biofilms in general, which play a crucial role in human health and disease.

## Methods

### Cultivation of macrocolony biofilms

In this study, we used the uropathogenic strain *E. coli* UTI89, which is frequently utilized as an *in vitro* biofilm model for investigating urinary tract infections (Song et al., 2022; Hadjifrangiskou et al., 2012; Beebout et al., 2019; García et al., 2021). We obtained this strain, which was originally isolated from a patient with an acute bladder infection (Chen et al., 2006), from the Danish Technical University, Biosustain.

The starting liquid culture (LC) was prepared by incubating the strain in liquid LB medium (1% Bacto tryptone, 0.5% Bacto yeast extract, 0.5% NaCl) for 16 h at 37 °C. Macrocolony biofilms were grown as described in the optimized protocol outlined in Serra and Hennge (2017), using an alternative YESCA agar medium (1% casamino acids, 0.1% yeast extract, 2.0% agar). The agar plates were incubated at 27 °C and the entire macrocolony biofilms were harvested for RNA and protein extraction at 6 and 12 hours, and at 1, 1.5, 2, 3, 5, 7, 14, 21, and 30 days post-inoculation (Figure 1a). Biofilm samples were labelled as 6H, 12H, 1D, 1.5D, 2D, 3D, 5D, 7D, 14D, 21D, 1M respectively. All samples were taken in three biological replicates per time-point.

Biofilm samples of spatial regions were collected at 1D (early-stage), 3D (mid-stage), and 14D (late-stage) stages of biofilm growth. Using a scalpel, we dissected biofilms at 1D and 3D stages into two parts (inner, outer) and at 14D stage into three parts (inner, middle, outer) (Figure 4a). Spatial regions from several biofilms were pooled per sample to reach a sufficient amount of bacterial material for RNA and protein extraction. The samples were taken in three biological replicates per spatial region.

### Imaging of macrocolony biofilms

A time-lapse video was created to monitor the growth of biofilms, using Mirrorless SONY alpha 7 II camera. The camera was coupled with a 144 LED light-ring mounted on a Zeiss Stemi C-2000 stereo microscope using a T2 adapter. This setup facilitated the observation of biofilm morphology development and was used to determine sampling time-points (Futo et al., 2022). For the time lapse video recording, biofilms were incubated at 27 °C with an average relative humidity of 80% using the automatic camera settings to ensure consistent recording conditions (Futo et al., 2022). The time lapse video was generated from 1,187 shots captured over a period of 25 days, with each shot taken at 30 min intervals. Adobe After Effects CC 2017 software was utilized to compile the shots into a video format, adjusting the frame rate to 24 fps.

### RNA extraction

The starting liquid culture (LC) of *E. coli* UTI89 grown in LB medium was centrifuged for 10 min 5,000 x *g* before RNA extraction. A scalpel and the upper parts of pipette tips were used to section biofilms regions (inner, middle, outer region) for spatial biofilm sampling. Entire biofilms and their spatial regions were harvested from agar plates with plastic spreaders and transferred into 2 mL tubes with 1.5 mL of stabilization buffer, RNAprotect Bacteria Reagent (Qiagen) diluted with PBS in a 2:1 volume ratio.

Total RNA was extracted using a modified version of the RNeasy Protect Mini Kit protocol (Qiagen). Disruption and homogenization of biofilm samples was done with a rotor-stator homogenizer Tissue Ruptor (Qiagen) at level 3 for 10 s with a 7 mm probe. Homogenized samples were vortexed for 10 s and incubated for 5 min at room temperature (RT), and then divided into three new 2.0 mL tubes for lysis. After centrifugation for 10 min at 5,000 x *g* at RT, the supernatants were discarded. Pellets were resuspended with 220 µL of the mix containing 200 µL of TE buffer (30 mM TrisHCl, 1 mM EDTA) with lysozyme (15 mg/mL) and 20 µL of Proteinase K (200 mg/mL). During lysis, samples were incubated in a shaker for 20 min at 25 °C and 550 rpm.

Following this incubation step, 3 volumes of RLT buffer (Qiagen) per tube were added and the mixture was vortexed vigorously. After centrifugation for 10 min at 8,000 x *g*, the supernatants were transferred into new 2.0 mL tubes, and 0.5 volumes of chloroform was added. After 10 min incubation at RT, samples were centrifuged for 15 min at 13,000 x *g* at 4 °C. The upper phase of centrifuged samples was pipetted into new tubes and 1 volume of 80% ethanol was added to all samples. Each suspension was gently mixed by pipetting, transferred into a mini spin column (Qiagen), and centrifuged for 15 s at 10,000 x *g*. Flow-through was discarded and the step was repeated until all suspension was centrifuged through the column. The columns were then washed with 700 µL of RW1 buffer (Qiagen) and then centrifugated for 15 s at 10,000 x *g*. The flow-through was discarded and the washing step with RPE buffer (Qiagen) was repeated two times with centrifugation for 1 min at 10,000 x *g*. The collection tubes were replaced with new 1.5 mL tubes, and 50 µL of RNase-free water was added to the column membrane, followed by a 1-minute incubation at room temperature (RT). Total RNA was then eluted from the column membrane by centrifugation for 1 minute at 10,000 x g.

In the next step, DNase I stock solution (Qiagen) was mixed with RDD buffer (Qiagen), added to samples, and incubated for 10 min at RT. After DNA digestion, LiCl with final concentration 3.75 M was added to samples and the mixture was incubated over-night at -20 °C. After an over-night incubation, samples were centrifuged for 15 min at 16,000 x *g* at 4 °C and rinsed with 80% ethanol. Supernatants were discarded and the pellets were dried before addition of 30 µL RNase-free water. Samples were kept on ice during the RNA quantification on NanoDrop 2000 spectrophotometer (Thermo Fisher Scientific) and during analysis of RNA integrity on agarose gel-electrophoresis. The RNA samples were then stored at -80 °C until sequencing.

### RNA sequencing

For total RNA sequencing, a combined approach was used, involving rRNA depletion followed by library preparation using the TruSeq Stranded kit with the NEB rRNA Depletion Kit (Illumina). RNA sequencing was performed bi-directionally on the Illumina NovaSeq 6000 S4 platform at Macrogen Europe BV (Maastricht, the Netherlands), generating between 65 to 90 million reads per run for both entire and spatial biofilm samples.

### Transcriptome data analyses

Before mapping, the sequence quality was checked using FastQC v0.11.9 (Andrews et al., 2010). In average, 86,846,330 paired-end sequences (100 bp read length) per sample were mapped onto the *E. coli* UTI89 reference genome (NCBI Assembly accession: ASM1326v1; GCA_000013265.1) using BBMap v38.39 (Bushnel, 2014) with an average of 98.3% mapped reads per biofilm sample (Supplementary File S1). One of the biological replicates for the 21D time-point failed during RNA sequencing library preparation and was thus excluded from further analysis. Similarly, one biological replicate for the liquid culture (LC) time-point was excluded due to a low number of counts. Therefore, for the LC and 21D time points, we used two biological replicates instead of three. For spatial biofilm samples, on average 76,080,157 paired-end sequences (100 bp read length) per sample were mapped onto the same reference genome with an average of 99.6% mapped reads per sample (Supplementary File S15). The mapping was performed using default settings, with the option to trim the read names after the first whitespace enabled.

SAMtools package v1.9 (Li et al., 2009) was employed to generate, sort and index BAM files for downstream data analysis. RNAseq data was then processed in R v4.2.2 (R Development Core Team 2008) using custom-made scripts published in Futo et al. (2021). Furthermore, mapped reads were quantified per each *E. coli* UTI89 open reading frame using the rsamtools R package v2.14.0 (Morgan et al., 2017). Raw counts for 5,087 open reading frames (ORFs) in entire biofilm data and raw counts for 5,081 ORFs in spatial biofilm data were obtained using the GenomicAlignments R package V1.34.1 (Lawrence et al., 2013) (Supplementary File S1, Supplementary File S15).

Differential expression across time-points or between biofilm regions and replicates was assessed using PCA implemented in the DESeq2 V1.38.3 R package (Love et al., 2014) and visualized in the R ggplot2 package V3.5.0 (Wickham, 2016) (Figure 1bd, Figure 4bd). The raw counts in both datasets (both entire and spatial biofilm data) were normalized by calculating the fraction of transcripts (τ) (Li et al., 2010). Following the approach in Futo et al. (2021) paper, the multiplication step was skipped and τ values were directly utilized for downstream calculations of evolutionary measures. Normalized replicates were resolved by calculating their median and excluding those with zero raw values. The resulting normalized transcript expression values were further used in phylo-transcriptomic analyses (see Phylostratigraphic analysis).

For RNA expression profile correlation, and visualization, genes with zero expression values across multiple samples in both datasets were discarded, reducing the dataset to 4,961 genes for the entire biofilm dataset and 4,989 genes for the spatial biofilm dataset. If a gene exhibited zero expression value in only one stage, the missing value was interpolated using the mean of two neighboring stages. If a zero-expression value occurred at the first or last stage of biofilm ontogeny, it was replaced directly with the value of the adjacent stage (Futo et al. 2021). To bring gene expression profiles to the same scale, each gene underwent normalization to the median and log2 transformation, resulting in per-gene standardized expression values (Supplementary File S2, Supplementary File S12, Supplementary File S14, Supplementary File S28). Visualization was done using the R ggplot2 package v3.5.0. Pearson’s correlation coefficients (R) were calculated for all-against-all comparison and represented in heatmaps, to assess transcriptome similarity across biofilm ontogeny stages (Figure 1ce) and regions (Figure 4ce). We utilized the DESeq2 V 1.38.3 (Love et al., 2014) R package’s pipeline to estimate the overall differential expression of each gene across all biofilm developmental stages and spatial regions within biofilm, employing the likelihood ratio test (LRT) (Supplementary File S16, Supplementary File S24).

### Protein extraction

The liquid culture (LC) of *E. coli* UTI89 grown in liquid LB medium was centrifuged for 10 min at 5,000 x *g* prior to protein extraction. Entire and spatial biofilm samples from agar plates were harvested in the same manner as for RNA extraction using plastic spreaders and transferred into 2.0 mL tubes with 1 mL of cell lysis buffer (4% w/v SDS, 100 mM Tris pH 8.5, 5 mM beta-glycerophosphate, 5 mM NaF, 5 mM Na3VO4, 10mM EDTA, and 1/10 tablet of Mini EDTA-free Protease Inhibitor Cocktail from Sigma Aldrich). Samples were resuspended in the cell lysis buffer and boiled for 10 min at 95 °C. Additional lysis was achieved by sonication on ice at 40% amplitude with 10 pulses repeated three times using the homogenizer Ultrasonic processor (Cole Parmer).

Debris was pelleted by centrifugation for 30 min at 10,000 x *g* at 4 °C. Soluble proteins from the supernatant were precipitated through chloroform-methanol precipitation. Four volumes of 99.99% methanol, one volume of chloroform, and three volumes of milliQ water were added to samples, followed by vortexing and centrifugation for 10 min at 5,000 x *g* at 4 °C. The upper aqueous phase was discarded without disturbing the protein-containing interphase. An additional four volumes of methanol were added to the tube, followed by vortexing, and centrifugation for 10 min at 5,000 x *g* and 4 °C. The supernatant was discarded, and the pellet was air-dried for 10 min. The air-dried pellet was dissolved in denaturation buffer (6 M urea, 2 M thiourea, 10 mM TrisHCl pH 8.0).

Protein concentration of the samples was estimated by Bradford method. 50 µg of total protein per sample was reduced with 20 mM dithiothreitol (DTT) for 30 min at room temperature. Reduced protein samples were then alkylated with 40 mM iodoacetamide (IAA) for 30 min at room temperature and protected from light. The alkylation reaction was quenched by adding a final concentration of 10 mM DTT. Protein samples were dissolved in 50 mM ammonium bicarbonate buffer pH 7.5 (ABC) to dilute the remaining urea in the samples to less than 1M urea. Protein digestion was performed in-solution by adding 1 µg of Pierce^TM^ Trypsin/Lys-C Protease solution (Thermo Scientific) to samples and incubation over-night at 37 °C.

Before mass spectrometry analysis, the resulting peptides were purified using SOLAμ™ SPE cartridges (Thermo Scientific). The column was spun through at 1,500 rpm for 1 min between each load. Initially, the columns were activated with 200 µL 100% methanol and then equilibrated with a load of 200 µL of 80% acetonitrile (ACN), 0.1% formic acid (FA), followed by two loads of 200 µL of 3% ACN, 1% triflouraceticacid (TFA). Samples were loaded onto the column and washed with two loads of 200 µL of 0.1% FA. Peptides were eluted twice with 30 µL of 40% ACN, 0.1% FA, dried in a vacuum centrifuge, and then analyzed using the mass spectrometry.

### Mass spectrometry

The mass spectrometry analyses were performed at The Proteomics Core Facility at Technical University of Denmark. Dried peptides were resuspended in 0.1% FA and loaded onto a 2 cm C18 trap column (ThermoFisher 164946), connected in-line to a 15 cm C18 reverse-phase analytical column (Thermo EasySpray ES904) using 100% of 0.1% FA in water at 750 bar, using the Thermo EasyLC 1200 HPLC system, and the column oven operating at 30 °C. Peptides were eluted over a 140 min gradient ranging from 6% to 60% of 80% ACN, 0.1% FA at 250 nL/min, and the Exploris 480 instrument (Thermo Fisher Scientific) with FAIMS Pro^TM^ Interface (ThermoFisher Scientific) switched between compensation voltage of -50 V and -70 V with cycles of 38 and 28 scans. Full MS spectra were collected at a resolution of 60,000, with an automatic gain control (AGC) target of 300%, maximum injection time set to auto, and a scan range of 375 – 1,500 m/z. MS1 precursors with an intensity of > 5×10^3^ and charge state of 2 to 6 were selected for MS2 analysis. Dynamic exclusion was set to 60 s, the exclusion list was shared between CV values and Advanced Peak Determination was set to ‘on’. Precursors selected for MS2 were isolated in the quadrupole with a 1.6 m/z window. Ions were collected with maximum injection time set to auto and normalized AGC target set to 75%.

Fragmentation was performed with an HCD normalized collision energy of 28% and MS2 at a resolution of 15,000. The MaxQuant software suite v2.4.2.0 (Cox and Mann, 2008; Tyanova et al., 2016) integrated with an Andromeda (Cox et al., 2011) search engine was utilized for the processing of acquired MS spectra. Database searching was conducted against a target-decoy database of *E. coli* UTI89 (NCBI Assembly accession: ASM1326v1; GCA_000013265.1) which contains 5,211 protein entries. The protease setting was specified as Endoprotease Trypsin/P, with a maximum missed cleavage allowance of two. Carbamidomethylation (Cys) was setup as a fixed modification. Label-free quantification was applied with a minimum ratio count of two. A false discovery rate (FDR) of 1% was applied individually at the peptide and protein levels for identification filtering with initial mass tolerance set to 20 ppm. Intensity-based absolute quantitation (iBAQ) option was enabled with log fit deactivated. All other parameters remained at their default settings. Subsequently, iBAQ values were obtained for 2,525 protein entries in the entire biofilm data and for 2,355 protein entries in the spatial biofilm data. In the entire biofilm data, the third biological replicate for the 12H replicate was excluded from the dataset due to the low number of predicted proteins.

### Proteome data analyses

Expression similarity across time-points and replicates for 2,518 proteins, as well as expression similarity across regions of biofilms and replicates for 2,355 proteins was evaluated using Principal component analysis (PCA) stats package in R v4.2.2 (R Development Core Team 2008). The PCA plot was generated using the R ggplot2 package v3.5.0 (Wickham, 2016) (Figure 1d, Figure 4d), implementing raw IBAQ values (Supplementary File S1, Supplementary File S15). Normalization of proteome data was done by dividing each iBAQ value by sum of all iBAQ values in the sample to calculate partial concentrations in the entire biofilm dataset and in the spatial biofilm dataset. Replicates were resolved by calculating their median, excluding replicates with zero iBAQ values. This process yielded normalized protein expression values, which were then used to compute evolutionary measures (see Phylostratigraphic analysis).

In proteome data for spatial samples, the third biological replicate for 14D and 3D time-points of inner biofilm region is missing due to uncharacteristically low number of predicted proteins. For preparation of the proteome data for protein expression profiles, transcriptome-proteome correlation and visualization, we eliminated genes with zero-expression values in more than one stage, resulting in 1,910 proteins for the entire biofilm dataset and 2,057 proteins for the spatial biofilm dataset. When a protein exhibited only one stage with a zero-expression value, we interpolated it by taking the mean values of the two neighboring stages. If a zero-expression value occurred in the first or last stage of the biofilm ontogeny, we directly assigned it the value of the only neighbor. To bring the protein expression profiles to the same scale, we normalized each gene’s expression to the median and then performed a log2 transformation on the obtained values, resulting in per-gene normalized expression values (standardized expressions) (Supplementary File S11, Supplementary File S14, Supplementary File S19, Supplementary File S28). We calculated the Pearson’s correlation coefficient (R) for the matching 1,858 genes/proteins to determine correlations between normalized transcriptome and proteome expression values at each biofilm time-point (Supplementary Figure 5, Supplementary File S14). Pearson’s correlation coefficients (R) were also calculated for all-against-all comparison to assess proteome similarity across biofilm ontogeny stages (Figure 1d, e) and biofilm regions (Figure 4 d, e). To estimate the overall differential expression of each protein across all biofilm developmental stages and spatial regions within biofilm, we employing the likelihood ratio test (LRT) using the DESeq2 V 1.38.3 (Love et al., 2014) (Supplementary File S16, Supplementary File S26).

### Phylostratigraphic, phylotranscriptomic and phyloproteomic analyses

We implemented phylostratigraphic approach as previously described (Domazet-Lošo et al. 2007; Domazet-Lošo and Tautz, 2010a; Domazet-Lošo and Tautz, 2010b; Domazet-Lošo et al. 2017; Futo et al. 2021). Following the relevant phylogenetic literature (Zaremba-Niedzwiedzka et al. 2017; Mukherjee et al., 2017; Hug et al., 2016; Wu et al., 2009; Brandis, 2021; Brown and Volker, 2004; Sharma et al., 2022; Williams et al., 2010; Adelou et al., 2016; Janda et al., 2021; Dunne et al., 2017; Denamur et al., 2020; Yu et al., 2021), we constructed a consensus phylogeny covering the lineage from the last common ancestor of cellular organisms to our focal species, *E. coli UTI89* (Supplementary Figure 1, Supplementary File S21).

We retrieved the complete set of protein sequences for 922 terminal taxa present on our consensus phylogeny from the NCBI database. This collection of proteomes represents our reference protein sequence database. To construct the *E. coli* UTI89 phylostratigraphic map (Supplementary File S22) (Domazet-Lošo et al., 2007; Domazet-Lošo and Tautz, 2010a), we compared 5,211 *E. coli* protein coding genes to the reference database using the blastp algorithm v2.9.0 (Altschul at al., 1990) and the e-value threshold of 10^−3^. We assigned 5,206 protein sequences that passed the phylostratigraphic procedure to the nine phylostrata of the consensus phylogeny (Supplementary File S22) using the previously described pipeline (Futo et al., 2021). Each protein sequence was assigned to the oldest phylostratum in which it still had a blast hit (Domazet-Lošo et. al., 2007; Domazet-Lošo and Tautz, 2010b).

TAI is the weighted mean of phylogenetic ranks (phylostrata), whose statistical properties and the biological interpretation have been previously described (Domazet-Lošo and Tautz, 2010a):

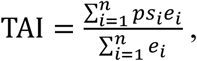

where *ps_i_* is an integer indicating the phylostratum of the protein *i*, *e_i_* is the normalized transcript expression value of the gene *i*, and *n* is the total number of genes analyzed. Using expression values for 4,984 protein coding genes we calculated the TAI for each developmental stage (entire biofilm dataset) using myTAI package v0.9.3. (Drost et al., 2018) (Figure 2a, Supplementary File S4). In the dataset for spatial biofilms, TAI was calculated using expression values of 4,979 protein coding genes (Supplementary File S18).

To analyze the proteome data in a similar way, we applied the PAI, *i.e.* the weighted mean of phylogenetic ranks (phylostrata):

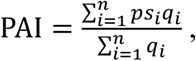

where *ps_i_* represents an integer-coded phylostratum assigned to protein *i*, *q_i_* is the normalized protein expression value of the protein *i*, and *n* corresponds to the total number of proteins analyzed. From the entire biofilms dataset, we used 2,518 proteins for PAI calculation and from spatial biofilms dataset, we obtained 2,355 proteins for PAI calculation (Figure 2b, Supplementary File S4, Supplementary File S18).

TAI and PAI profiles, which reflect the phylogeny-ontogeny correlation, were tested for statistical significance using the ‘Early conservation test’ (Drost et al., 2018). This test requires the predefinition of developmental modules corresponding to early, middle, and late biofilm stages. Based on PCA and Pearson’s correlation analyses of transcriptomic and proteomic data (Figure 1), we assigned developmental time points to modules as follows: the early stage from 6H to 1D, the mid stage from 1.5D to 3D, and the late stage from 5D to 1M.

Evolutionary divergence rates of *E. coli* UTI89 proteins were estimated with regard to *Klebsiella pneumoniaei* and *Yersinia pestis*. Comparison to these two bacterial species provided the best results regarding the number of detected orthologues and the suitability of evolutionary distances for calculating evolutionary rates. Following reciprocal best hits using blastp v2.13.0 with 10^−5^ e-value threshold, we obtained 3,137 orthologs in *K. pneumoniae* MGH78578 (NCBI Assembly accession: GCA_000016305.1; ASM1630v1) and 2,530 orthologs in *Y. pestis* biovar Microtus str. 91001 (NCBI Assembly accession: GCA_000007885.1; ASM788v1). We globally aligned *E. coli* UTI89 and each bacterial strain to obtain orthologous pairs using the Needleman–Wunsch algorithm and then constructed codon alignments in pal2nal (Suyama et al., 2006). The non-synonymous substitution rate (dN) and the synonymous substitution rate (dS) were calculated according to Comeron’s method (Comeron, 1995). The entire procedure to determine dN and dS was performed using R package orthologr v0.4.0 (Drost et al., 2015).

Using dN values calculated separately for each bacterial species, we calculated the transcriptome non-synonymous divergence index (TdNI, Figure 2c, Supplementary Figure 2a, Supplementary File S6, Supplementary File S7), *i.e.* the weighted mean of non-synonymous gene divergence:

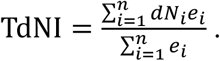

TdNI profiles are calculated using a real number representing the non-synonymous divergence of the gene *i* (d*N_i_*), normalized transcript expression value of the gene *i* (*e_i_*), and the total number of genes analyzed (*n*) (Futo et al., 2021).

Using dS values calculated separately for each bacterial strain, we calculated transcriptome synonymous divergence indices (TdSI; Supplementary Figure 2b, Supplementary Figure 3a, Supplementary File S6, Supplementary File S7), *i.e.,* the weighted mean of synonymous gene divergence:

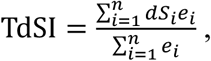

where d*S_i_* is a real number that represents the synonymous divergence of gene *i*, *e_i_* is the normalized transcript expression value of the gene *i*, and *n* is the total number of genes analysed (Futo et al., 2021).

To analyze proteome data in a similar way, we applied the PdNI (the proteome non-synonymous gene divergence index) and PdSI (the proteome synonymous gene divergence index), where normalized protein expression values were used instead of normalized transcript expression values as in TdNI and TdSI. Using dN and dS values of the above-mentioned bacterial strains, we calculated the proteome non-synonymous and synonymous divergence indices (PdNI, PdSI, Figure 2d, Supplementary Figure 2c,d, Supplementary Figure 3b, Supplementary File S6, Supplementary File S7), *i.e.,* the weighted mean of non-synonymous and synonymous divergence:

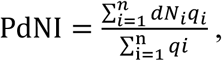

where d*Nᵢ* is a real number representing the nonsynonymous divergence of protein *i*, *qᵢ* denotes the normalized expression level of protein *i*, and *n* refers to the total number of proteins included in the analysis, and:

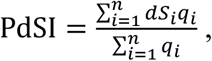

where d*Si* represents the synonymous divergence value of protein *i*, *q_i_* is the normalized protein expression value of protein *i* that acts as weight factor and *n* is the total number of proteins analyzed.

Transcriptome codon usage bias index (TCBI) is a weighted arithmetic mean calculated using 4,984 transcriptome expression values and measure independent of length and composition (MILC) (Supplementary Figure 3c, Supplementary File S5) (Supek and Vlahoviček, 2005; Futo et al., 2021):

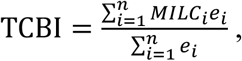

where MILC is a real number which represents the codon usage bias of gene *i*, *e_i_* is the normalized transcript expression value of the gene *i*, and *n* is the total number of genes analyzed. MILC value is a real number that represents the codon usage bias of a gene and was determined using R package coRdon v1.16.0 (Elek et al., 2019), in relation to codon usage bias of *E. coli* UTI89 ribosomal genes. Proteome codon usage bias index (PCBI) was calculated using 2,518 protein expression values and MILC values (Supplementary Figure 3d, Supplementary File S5) (Futo et al. 2021):

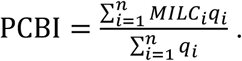

PCBI is weighted arithmetic mean of codon usage bias, where MILC is a real number which represents the codon usage bias of protein *i*, *e_i_* is the normalized protein expression value of the protein *i*, and *n* is the total number of proteins analyzed.

The statistical significance of difference in average TAI and PAI values between two biofilm regions (inner and outer) in 1D and 3D biofilms was assessed by t-test (Figure 5, Supplementary File S18). For 14-day-old biofilms, differences across three biofilm regions (inner, middle, and outer) were evaluated using ANOVA (Figure 5, Supplementary File S18).

### Enrichment analysis and functional annotation

Gene ontology (GO) functional annotations for the *E. coli* UTI89 strain were retrieved by searching for orthologs in the eggNOG v5.0 database (Huerta-Cepas et al., 2019) using the eggNOG-mapper v2.1.9 (Cantalapiedra et al., 2021). The best annotation data was obtained by using the default search filters and setting the taxonomic scope to Bacteria. This resulted in 3,108 genes with GO annotations. The names for GO annotation used for functional enrichments were downloaded from the Gene Ontology Resource website (https://geneontology.org/docs/download-ontology/, February 9, 2023). Synonyms and obsolete GO terms were cleaned to unify the GO terminology before enrichment analysis. Furthermore, custom annotations “GO:no_ID no annotation” were added to the genes that were not annotated by the eggNOG-mapper. To estimate functional enrichments of transcriptome and proteome data, we conducted one-tail hypergeometric tests to identify overrepresented functions. In all enrichment analyses, p-values were adjusted for multiple comparisons using the Benjamini and Hochberg procedure (Benjamini and Hochberg, 1995).

We analyzed functional enrichment GO annotations at individual developmental stages for transcriptome and proteome data (Figure 3, Supplementary Figure 6). Functional enrichment of GO categories was evaluated for upregulated genes with standardized expression ≥ 0.5 (log₂ scale) above the median and FC > 2 (Supplementary File S3), as well as for proteins with standardized expression ≥ 0.15 (log₂ scale) above the median and FC > 0 during biofilm development (Supplementary File S25). We also estimated the distribution of expressed functionally uncharacterized genes enriched across phylostrata during biofilm development (Supplementary Figure 4, Supplementary File S9).

Enrichment profiles of GO annotations for differentially expressed transcripts and proteins in individual biofilm regions. Pairwise differential expression analysis between inner, middle, and outer biofilm region of 1D, 3D, and 14D biofilms was estimated using DESeq2 package v1.38.3 (Love et al. 2014) (Supplementary File S23, Supplementary File S27). Functional enrichment of GO annotations was assessed for genes exhibiting a significant shift in expression (p-value < 0.05) with a fold change (FC) > 2, and for proteins with a significant shift in expression (p-value < 0.05) and a fold change (FC) > 0 (Supplementary Figures 7–10, Supplementary File S20, Supplementary File S25). We visualized the results of enrichment analyses using custom-made scripts based on the R ggplot2 package v3.5.0 (Wickham, 2016).

## Acknowledgments

This research was funded by the European Union’s Horizon 2020 research and innovation program under the Marie Skłodowska-Curie grant agreement 955626 (PEST-BIN).

## Author contributions

I.M., D.F., and T.D.-L. initiated and conceptualized the study, A.T. and M.F. grew the biofilms, performed wet lab experiments and visualized the biofilms, A.T., and E.S. quantified proteomes, A.T., S.K., M.F., N.K., N.Č., and M.D.-L. performed bioinformatic analyses, A.T., N.K., and T.D.-L. prepared the figures and tables for publication. A.T., N.K., and T.D.-L. wrote the manuscript. All authors read and approved the manuscript.

## Data availability

All transcriptome data have been deposited in NCBI’s Gene Expression Omnibus and are accessible through GEO Series accession number GSE272617. All mass spectrometry proteomics data have been deposited in the Proteome Xchange Consortium via the PRIDE partner repository with the dataset identifier PXD053727. The Supplementary File S13, contains a link to biofilm growth video available on YouTube: https://www.youtube.com/watch?v=rz--ezE-Wz4. All other data is provided within the manuscript or supplementary information files.

---

**Supplementary Figure 1.**
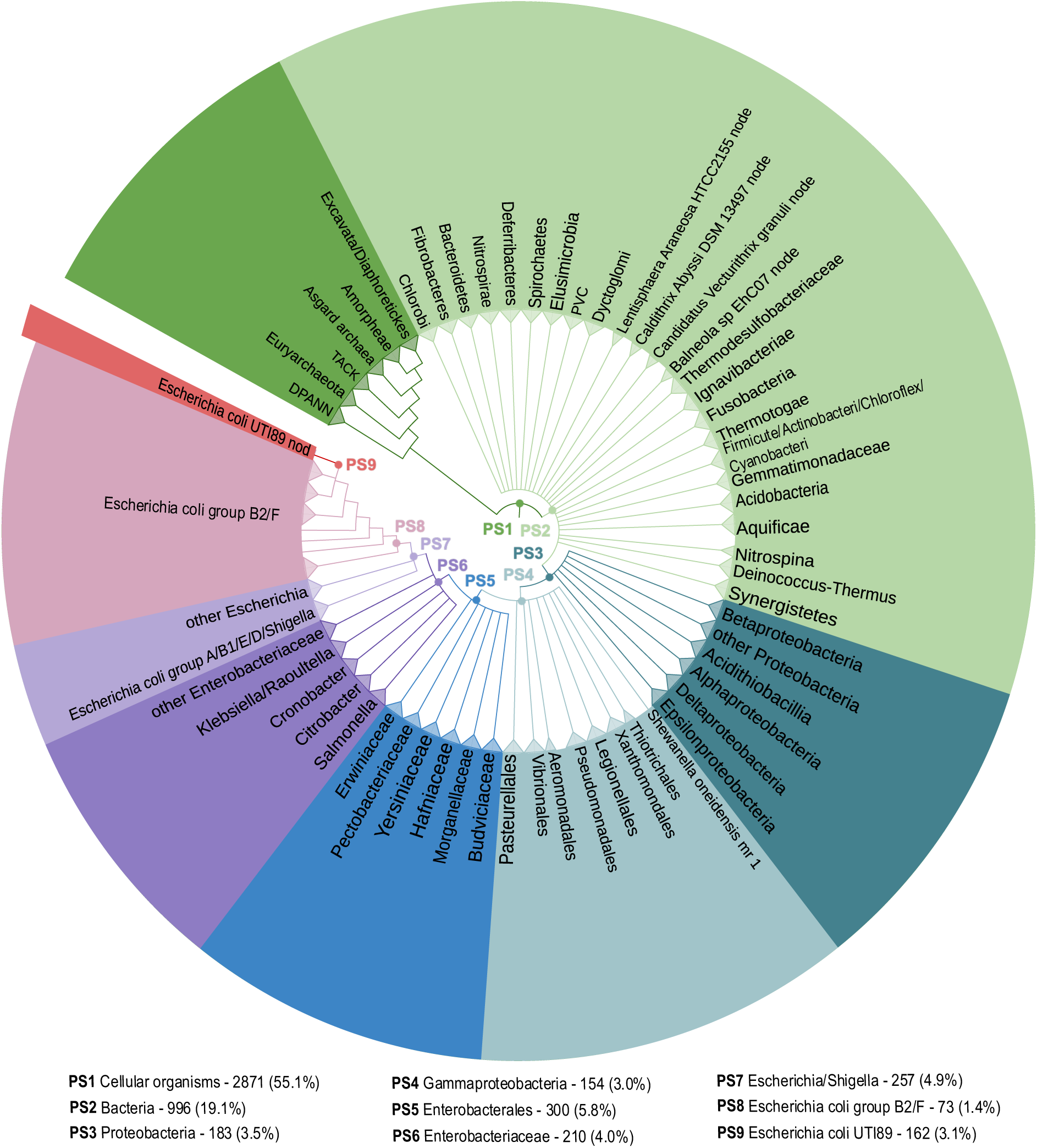
The consensus tree covers the divergence from the last common ancestor of cellular organisms to *E. coli* UTI89 as the focal species (see Supplementary File S21 for a fully resolved tree). The tree was constructed for phylostratigraphic analysis considering the evolutionary trajectory of focal strain and availability of reference genomes. The nine internodes (phylostrata) leading from the root of the tree to the focal species (*E. coli* UTI89) are labelled by ps1–ps9. The number of *E. coli* UTI89 genes that can be traced back to each phylostratum and the corresponding percentage are indicated after the name of the phylostratum.

**Supplementary Figure 2.**
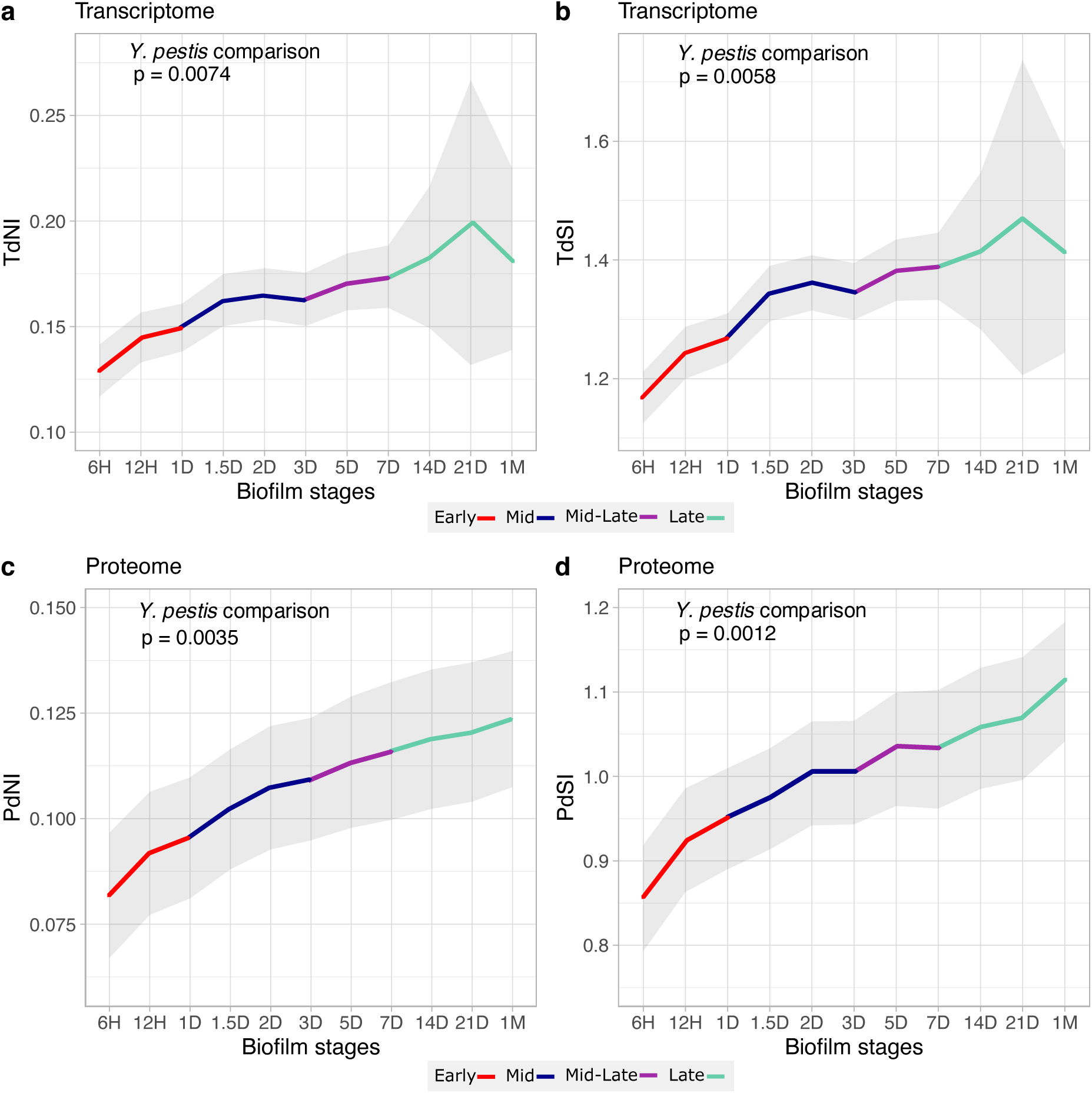
(**a**) Transcriptome non-synonymous divergence index (TdNI) and (**b**) transcriptome synonymous divergence index (TdSI) show an early conservation pattern. Synonymous and non-synonymous rates were estimated in *E. coli* UTI89 – *Y. pestis* pairwise comparisons. (**c**) Proteome non-synonymous divergence index (PdNI) shows early conservation pattern similar as (**d**) proteome synonymous divergence index (PdSI) (Supplementary File S7). The displayed p-values are derived from the Early conservation test, while the grey shaded regions indicate the range of ± one standard deviation estimated through permutation analysis. Biofilm growth stages, early (red), mid (blue), mid to late (purple), and late (green), are distinguished by color codes.

**Supplementary Figure 3.**
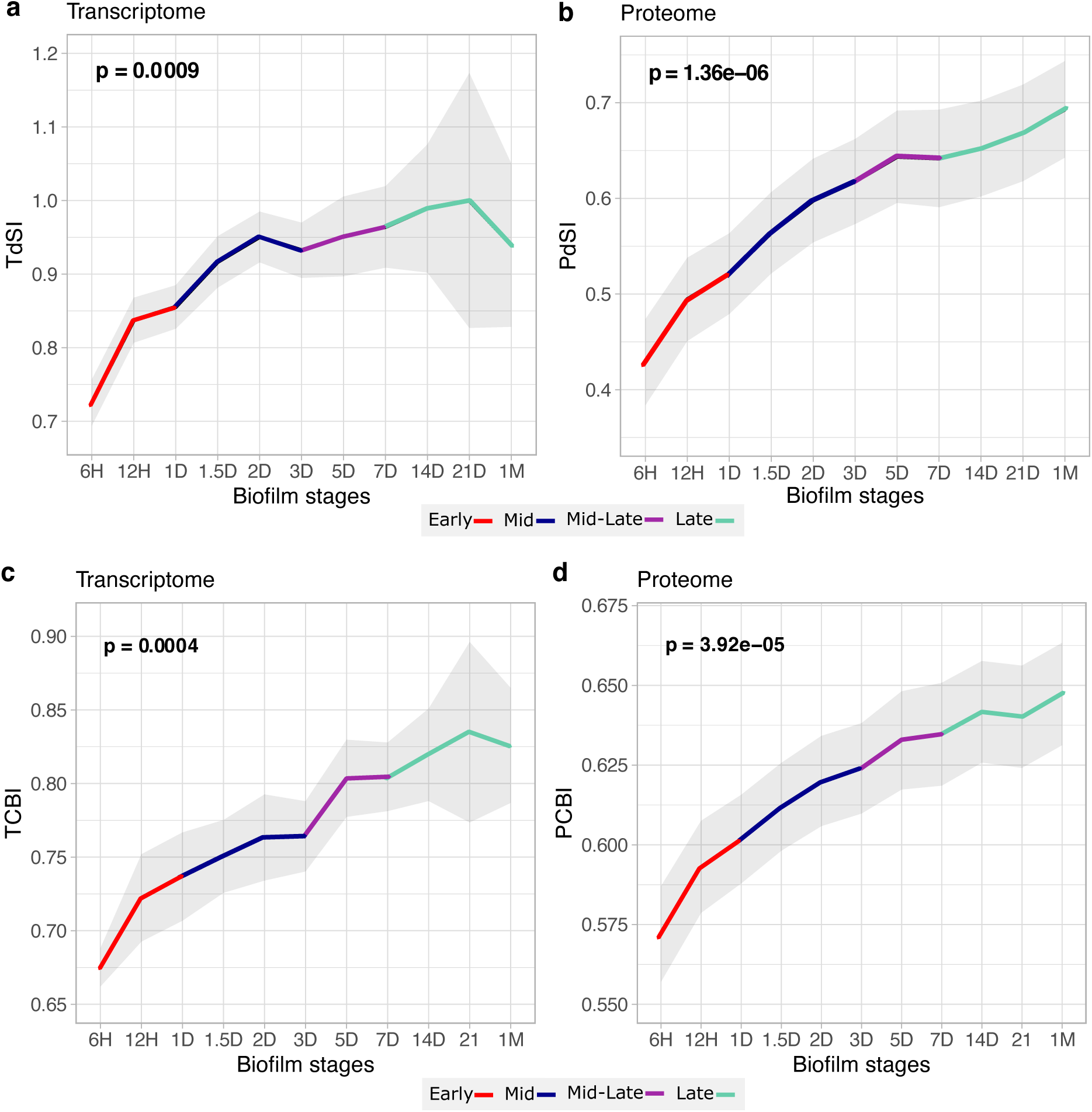
Ontogeny recapitulates phylogeny in uropathogenic *E. coli* biofilms through synonymous sites and codon usage bias. (**a**) Transcriptome synonymous divergence index (TdSI) and (**b**) proteome synonymous divergence index (PdSI) show an recapitulation pattern (Supplementary File S6). Early stages of biofilm ontogeny express conserved genes at synonymous divergence sites, while later stages express more diverged genes. Synonymous rates were estimated in *E. coli* UTI89 – *Klebsiella pneumoniae (*Enterobacteriaceae) comparisons. (Supplementary File S6). Pairwise comparison to additional *strain, Yersinia pestis* (Enterobacterales), showed similar results (Supplementary Figure 2, Supplementary File S7). Codon usage bias was calculated using MILC (see Methods). Both transcriptome codon bias index (TCBI) and proteome codon bias index (PCBI) show significant recapitulation pattern (Supplementary File S5). We evaluated significance (p-value) of these evolutionary patterns using early conservation test (see Methods). Grey shaded areas indicate ± one standard deviation based on permutation analysis. Different biofilm developmental stages are color-coded: early (red), mid (blue), mid-late (purple), and late (green).

**Supplementary Figure 4.**
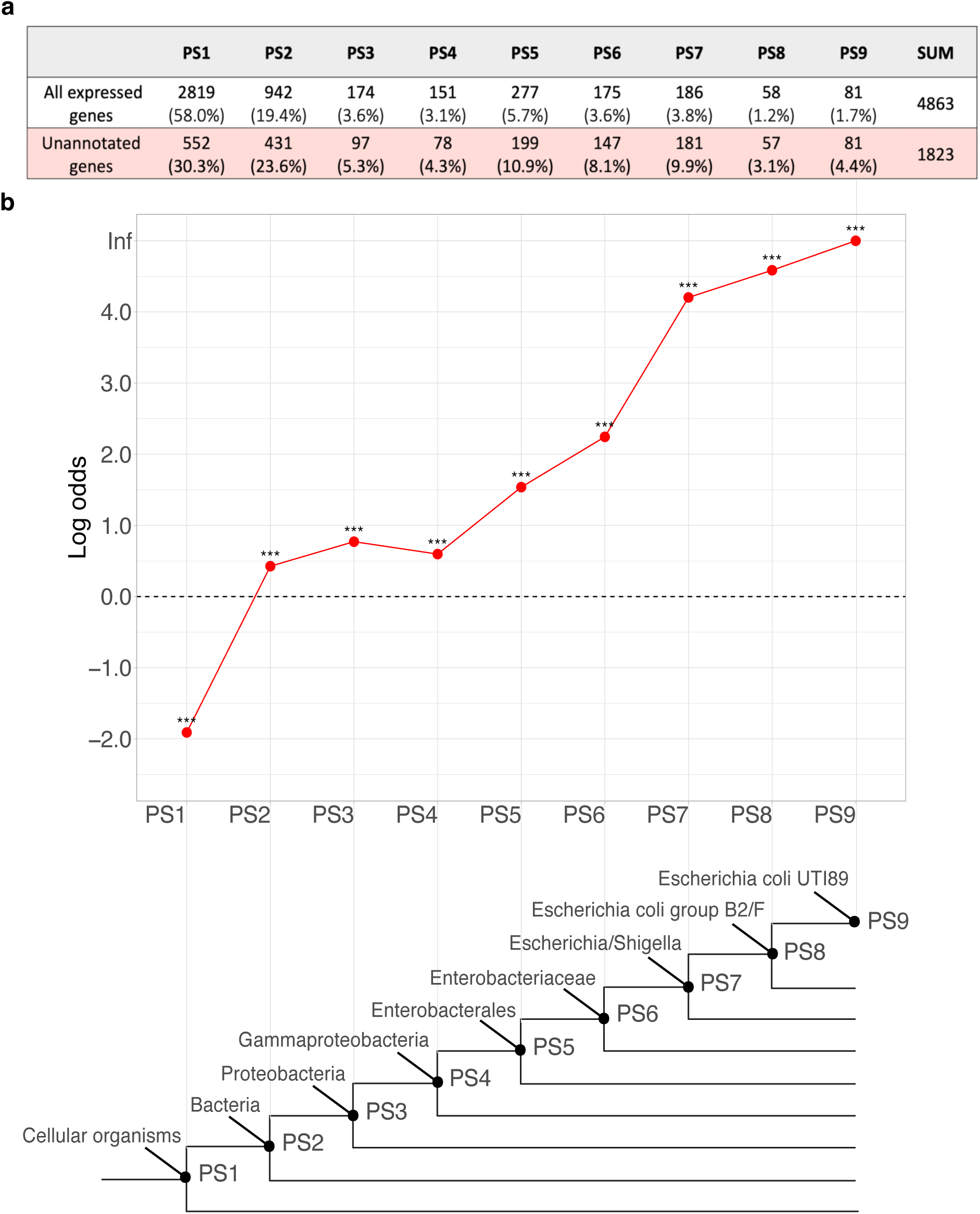
Evolutionary origin of functionally uncharacterized genes that are expressed during *E. coli* UTI89 biofilm development. (**a**) The table presents the number and percentage of expressed genes across phylostrata in two categories: all expressed genes and only those without functional annotation (unannotated genes). (**b**) Distribution of unannotated genes on the phylostratigraphic map. Statistical enrichment across phylostrata was assessed using a one-tailed hypergeometric test, with p-values adjusted for multiple testing (*** p < 0.001) (Supplementary File S9).

**Supplementary Figure 5.**
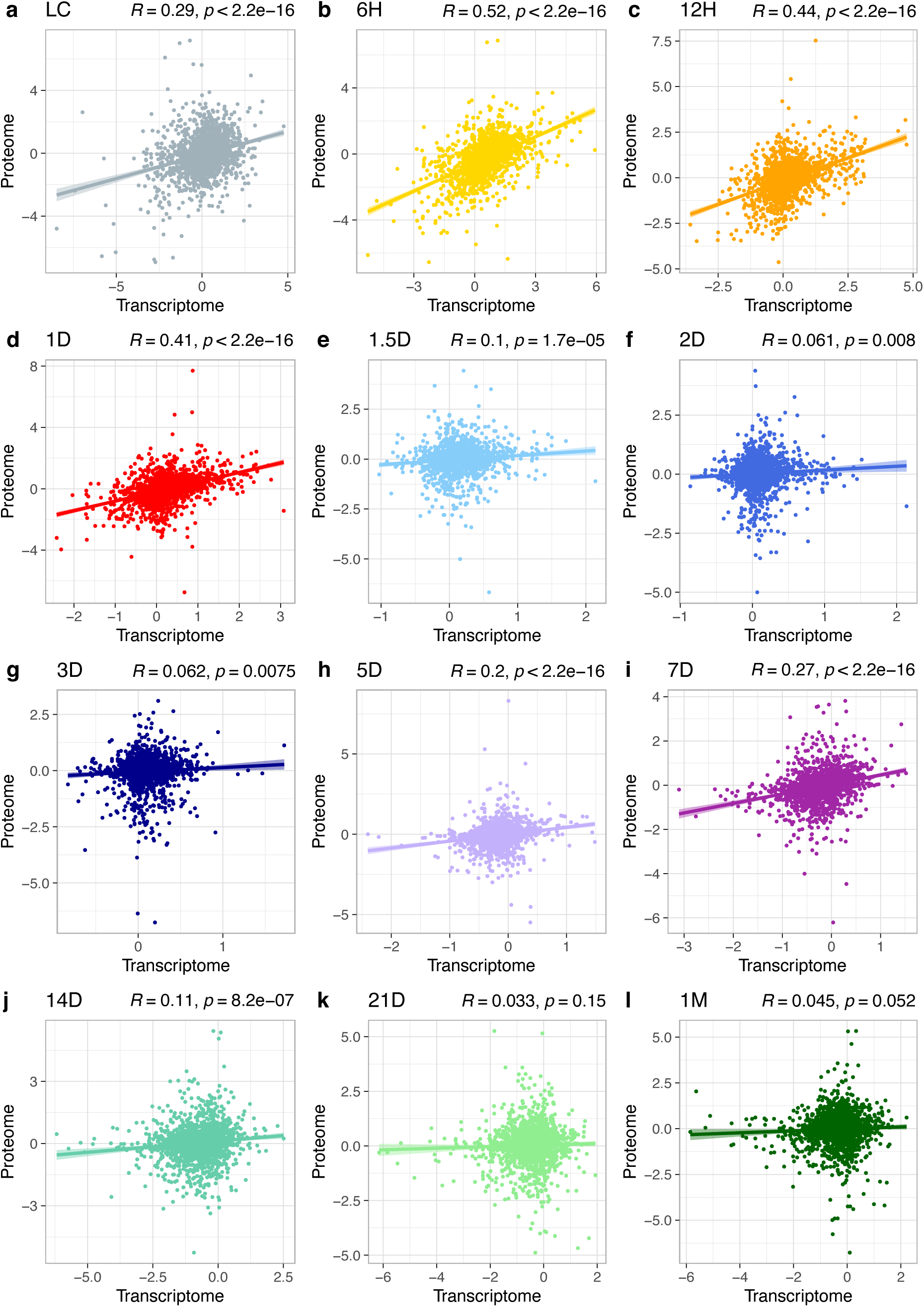
Transcriptome and proteome data overall show poor correlation, ranging from moderate (6H, 12H, 1D), low (LC, 5D, 7D), to very low (1.5D, 2D, 3D and 14D, 21D, 1M). Correlation analysis was conducted for all time-points during biofilm maturation, involving transcriptome normalized expression values on -log2 scale and proteome normalized expression values on -log2 scale (Supplementary File S14). Pearson’s correlation coefficients R with corresponding p-values are shown. Genes or proteins that had zero expression values in either transcriptome or proteome data were excluded.

**Supplementary Figure 6.**
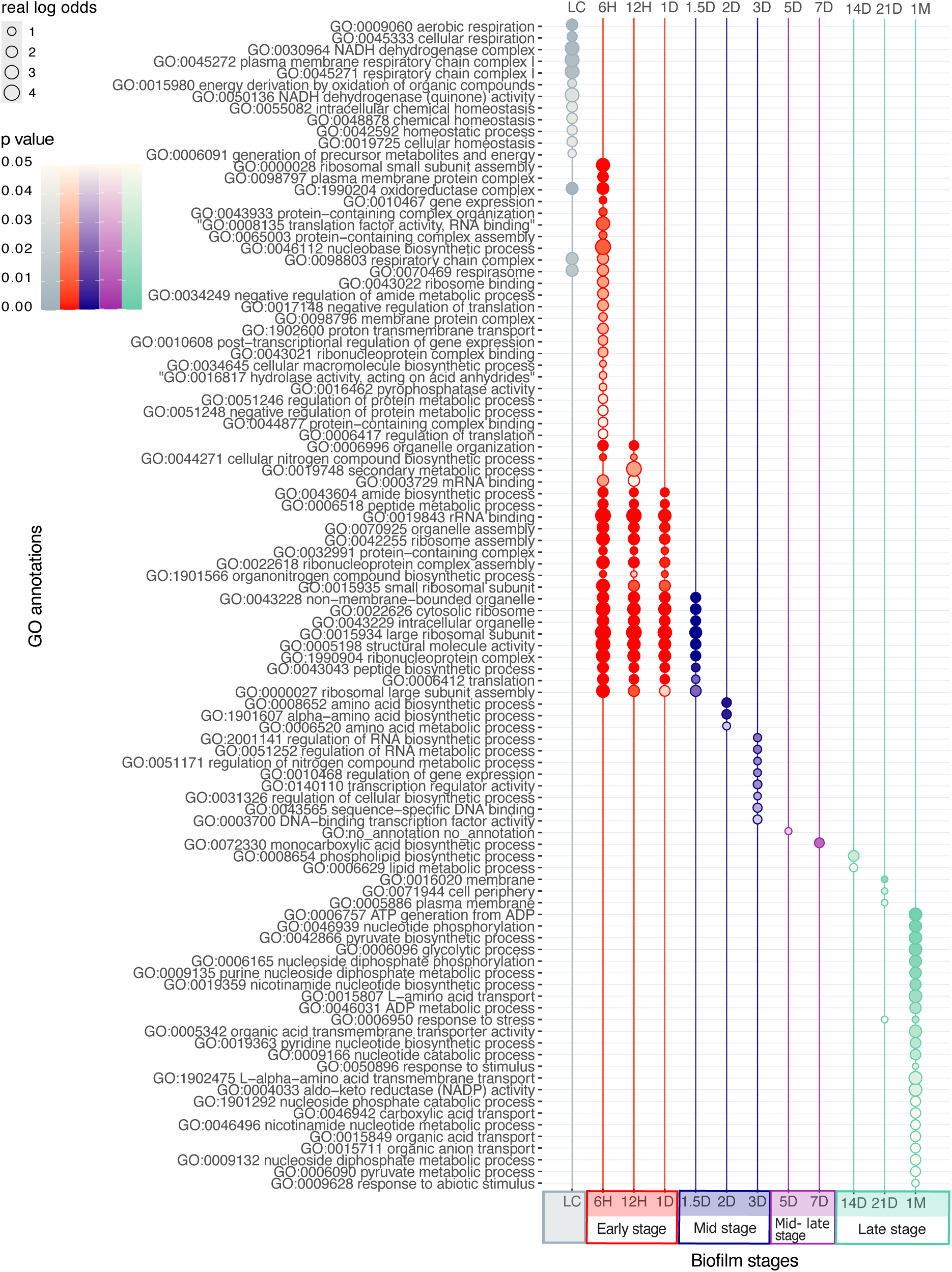
The temporal distribution of functional categories in the proteome data of *E. coli* UTI89 biofilms reveals distinct patterns. Functional enrichment of GO categories was assessed for upregulated proteins exhibiting expression levels at least 0.15 (log₂ scale) above the median during biofilm development. Enrichment significance was determined using a one-tailed hypergeometric test, with multiple comparisons corrected using the Benjamini-Hochberg procedure (significance threshold: p < 0.05; Supplementary File S25). Functional enrichment is visualized by real log odds (circle sizes), while significance is represented by color shades corresponding to p-values. Biofilm development stages are color-coded as follows: starting culture (grey), early stage (red), mid-stage (blue), mid-late stage (purple), and late stage (green).

**Supplementary Figure 7.**
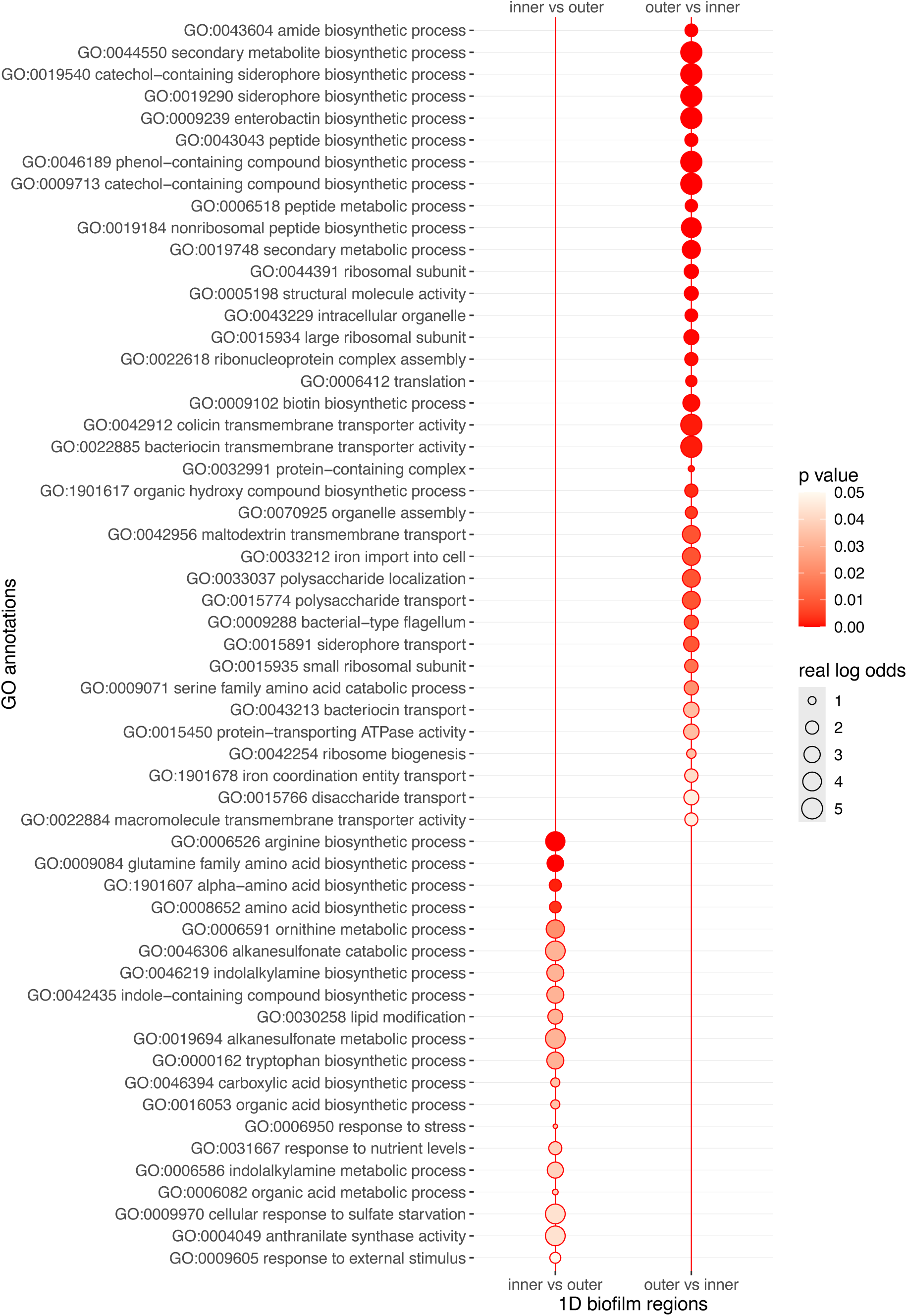
Spatially organized functions distribution in the early stage one-day-old (1D) biofilm. Enrichment profiles of GO functional annotations for upregulated genes that are differentially expressed genes in distinct parts of *E. coli* UTI89 biofilms. Differentially expressed genes from one 1D biofilm region were determined in reference to other 1D biofilm region using DeSeq2 pairwise comparison (Supplementary File S23). Genes were considered to be differentially expressed if the shift in its expression was statistically significant (p-value < 0.05) and additionally filtered for the magnitude of change to be at least two-fold (FC > 2). Significance of enrichment was tested using a one-tailed hypergeometric test for upregulated genes, corrected for multiple comparison at level below 0.05 (Supplementary File S20). The effect of functional enrichments is depicted by real log odds (circle sizes), while their significance is indicated by color shades corresponding to p-values.

**Supplementary Figure 8.**
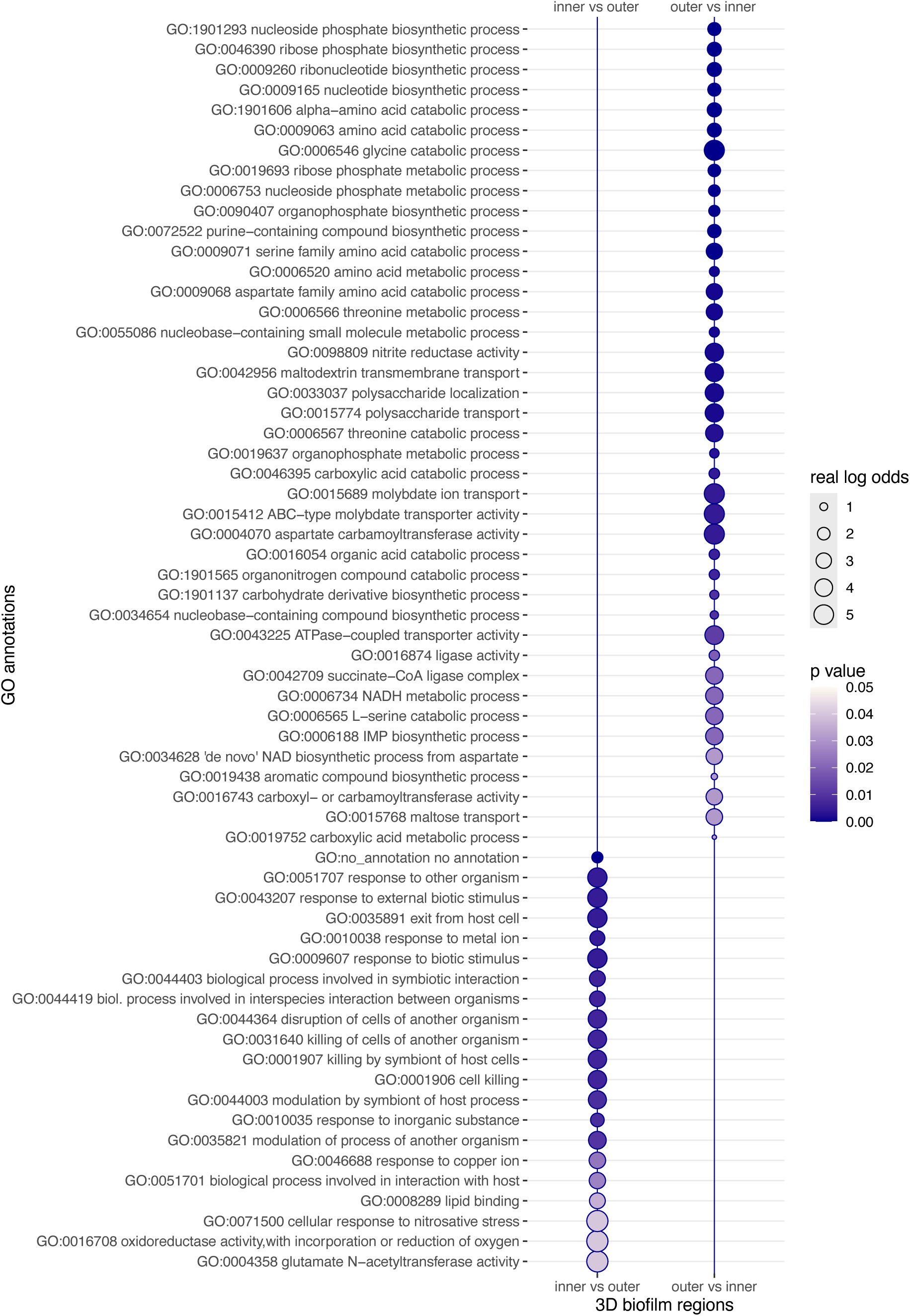
Spatially organized functions distribution in the mid stage three-day-old (3D) biofilm. Enrichment profiles of GO functional annotations for upregulated genes that are differentially expressed genes in distinct parts of *E. coli* UTI89 biofilms. Differentially expressed genes from one 3D biofilm region were determined in reference to other 3D biofilm region using DeSeq2 pairwise comparison (Supplementary File S23). Genes were considered to be differentially expressed if the shift in its expression was statistically significant (p-value < 0.05) and additionally filtered for the magnitude of change to be at least two-fold (FC > 2). Significance of enrichment was tested using a one-tailed hypergeometric test for upregulated genes, corrected for multiple comparison at level below 0.05 (Supplementary File S20). The effect of functional enrichments is depicted by real log odds (circle sizes), while their significance is indicated by color shades corresponding to p-values.

**Supplementary Figure 9.**
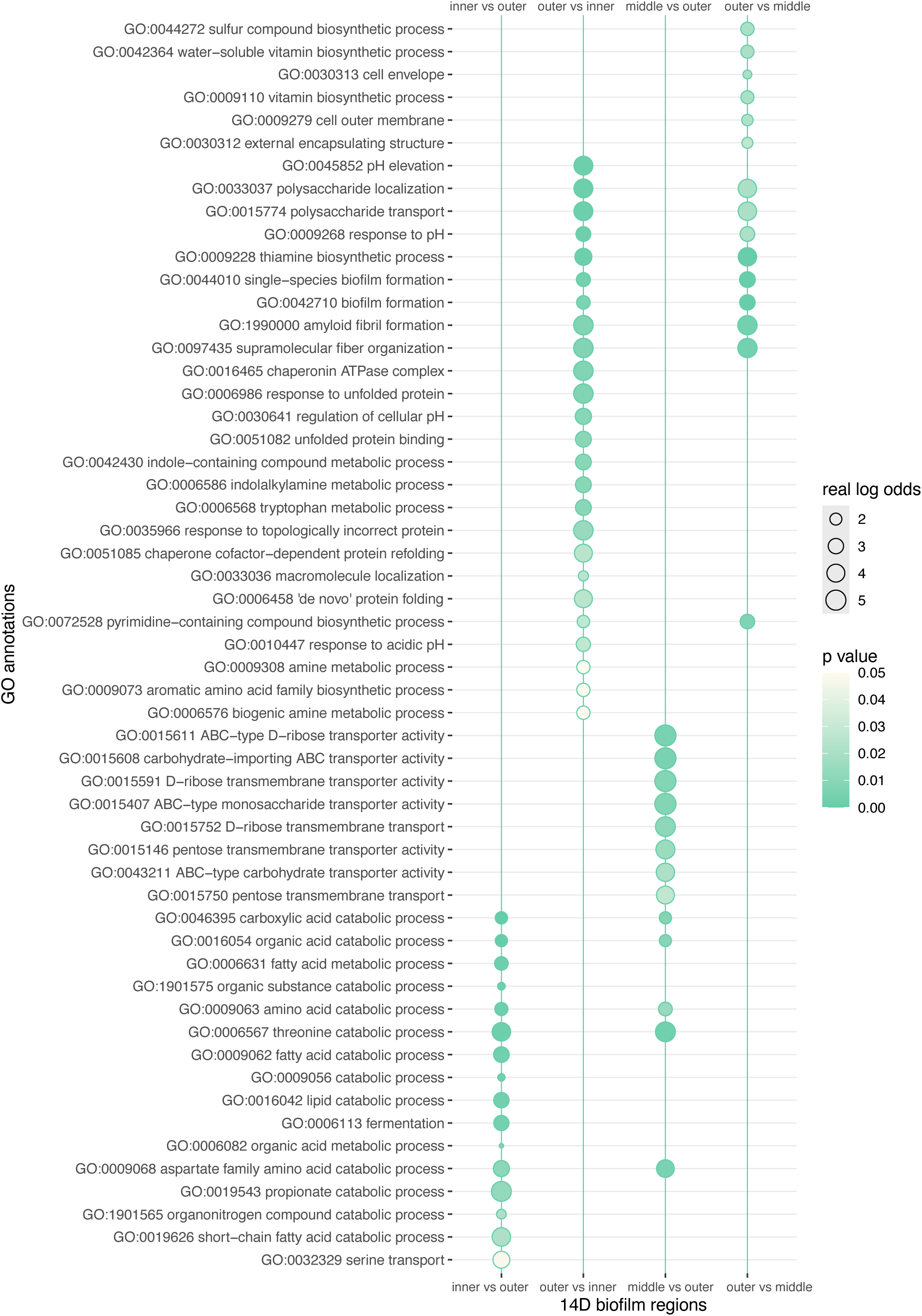
Spatially organized functions distribution in the late stage 14-day-old (14D) biofilm. Enrichment profiles of GO functional annotations for upregulated genes that are differentially expressed genes in distinct parts of *E. coli* UTI89 biofilms. Differentially expressed genes from one 14D biofilm region were determined in reference to other 14D biofilm region using DeSeq2 pairwise comparison (Supplementary File S23). Genes were considered to be differentially expressed if the shift in its expression was statistically significant (p-value < 0.05) and additionally filtered for the magnitude of change to be at least two-fold (FC > 2). Significance of enrichment was tested using a one-tailed hypergeometric test for upregulated genes, corrected for multiple comparison at level below 0.05 (Supplementary File S20). The effect of functional enrichments is depicted by real log odds (circle sizes), while their significance is indicated by color shades corresponding to p-values.

**Supplementary Figure 10.**
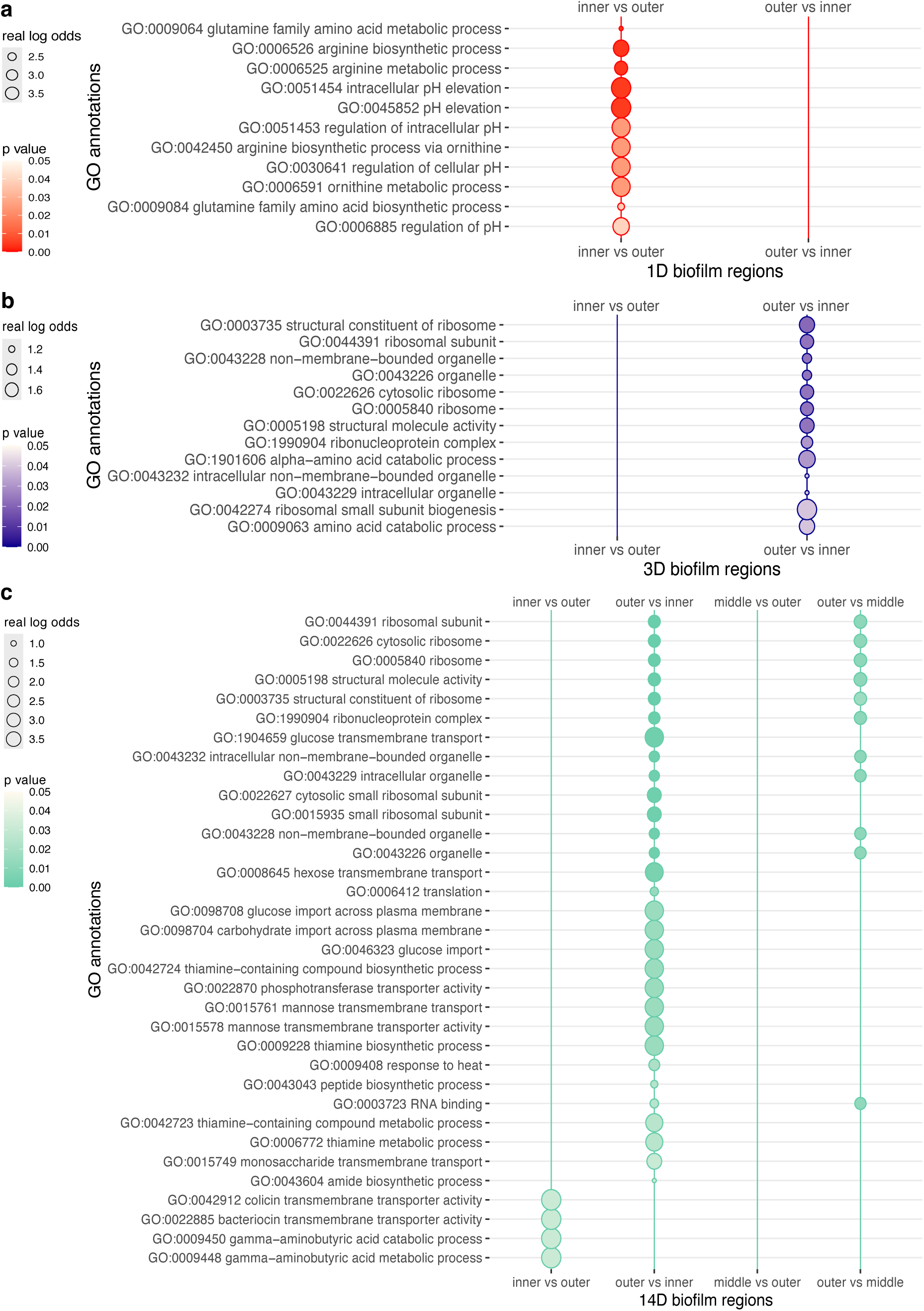
Spatially organized functions distribution in the proteome data of early stage (1D), mid stage (3D), and late stage (14D) *E. coli* biofilm development. Enrichment profiles of GO functional annotations for upregulated proteins that are differentially expressed genes in distinct regions of *E. coli* UTI89 biofilms. Differentially expressed proteins from one biofilm region were determined in reference to other biofilm region using DeSeq2 pairwise comparison (Supplementary File S27). Genes were considered to be differentially expressed if the shift in its expression was statistically significant (p-value < 0.05) and additionally filtered for the magnitude of change to be above 0 (FC > 0). Significance of enrichment was tested using a one-tailed hypergeometric test for upregulated genes, corrected for multiple comparison at level below 0.05 (Supplementary File S25). The effect of functional enrichments is depicted by real log odds (circle sizes), while their significance is indicated by color shades corresponding to p-values.

**Supplementary Figure 11.**
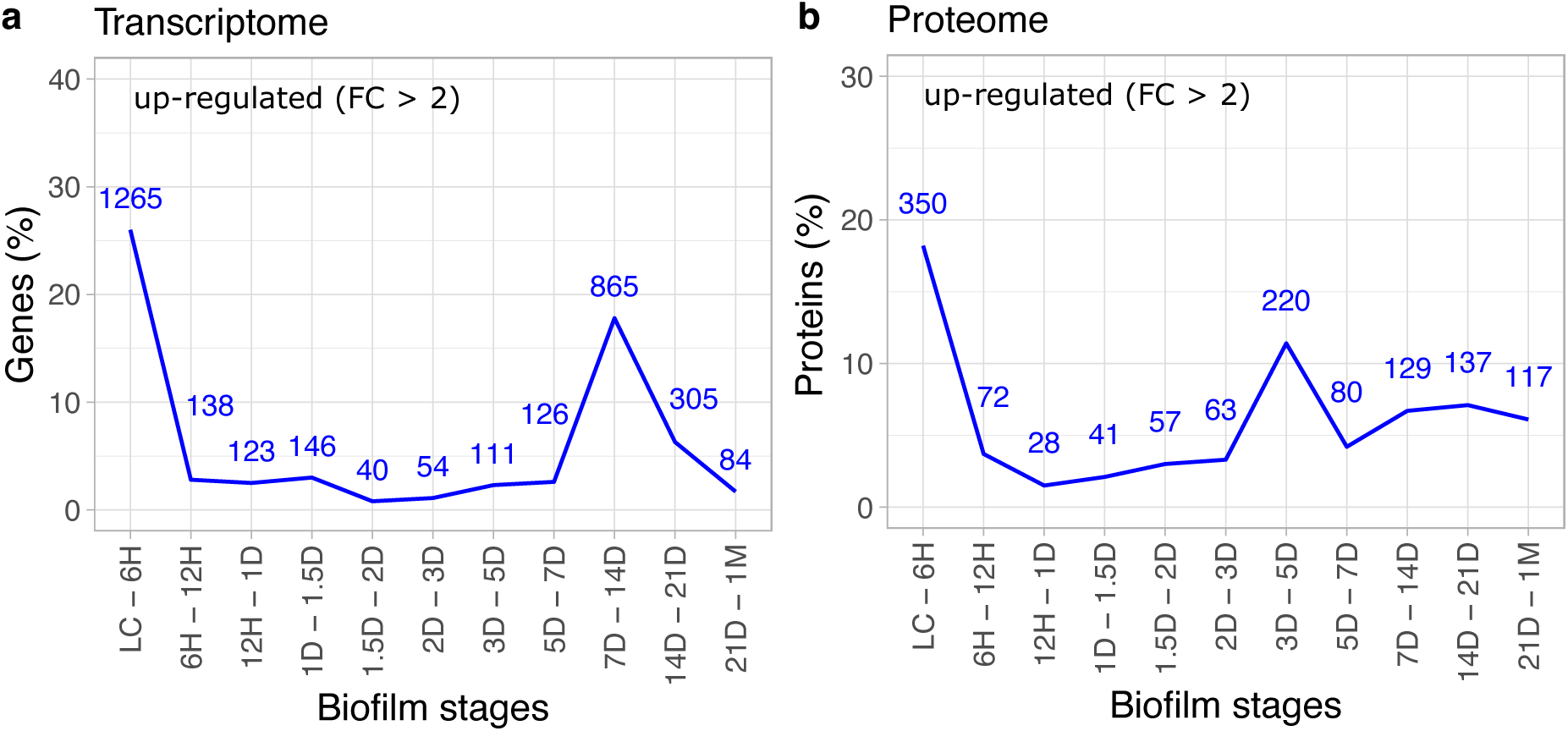
Differential expression in pairwise comparison between successive biofilm samples. Bursts of differentially expressed up-regulated genes are detected at inoculation of biofilm (LC – 6H) in both (**a**) transcriptome (Supplementary File S17) and (**b**) proteome datasets (Supplementary File S8). The numbers above each time-point indicate the number of differentially expressed genes or proteins. Pairwise differential expression was analyzed using DeSeq2, with p-values adjusted for multiple comparisons using the Benjamini and Hochberg procedure and cut off set at 0.05. Up-regulated (**a**) genes and (**b**) proteins are defined by a fold-change greater than 2.

**Supplementary Figure 12.**
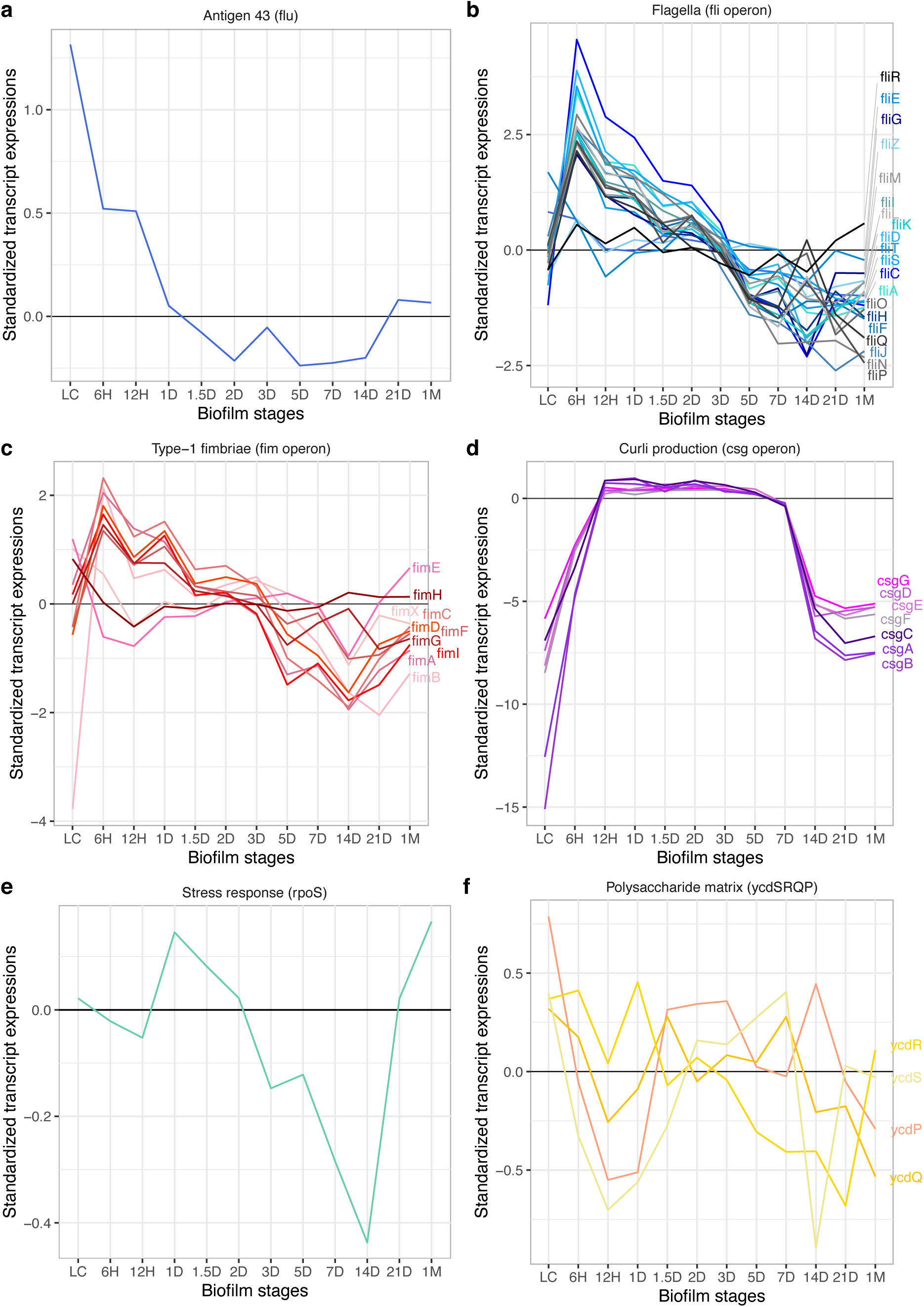
Standardized transcription profiles of key genes involved in rosette-initiated biofilm development: (**a**) Antigen 43 (*flu*) (n = 1), (**b**) flagella (*fli* operon) (n = 20), (**c**) type-1 fimbriae (*fim* operon) (n = 10), (**d**) curli production (*csg* operon) (n = 7), (**e**) stress response (*rpoS*) (n = 1), and (**f**) polysaccharide matrix (*ycdSRQP* operon) (n = 4). Grey horizontal line represents the median of standardized transcriptome expression values. For a full list of genes used in profile visualization and their corresponding expression values, refer to Supplementary File S14.

**Supplementary Figure 13.**
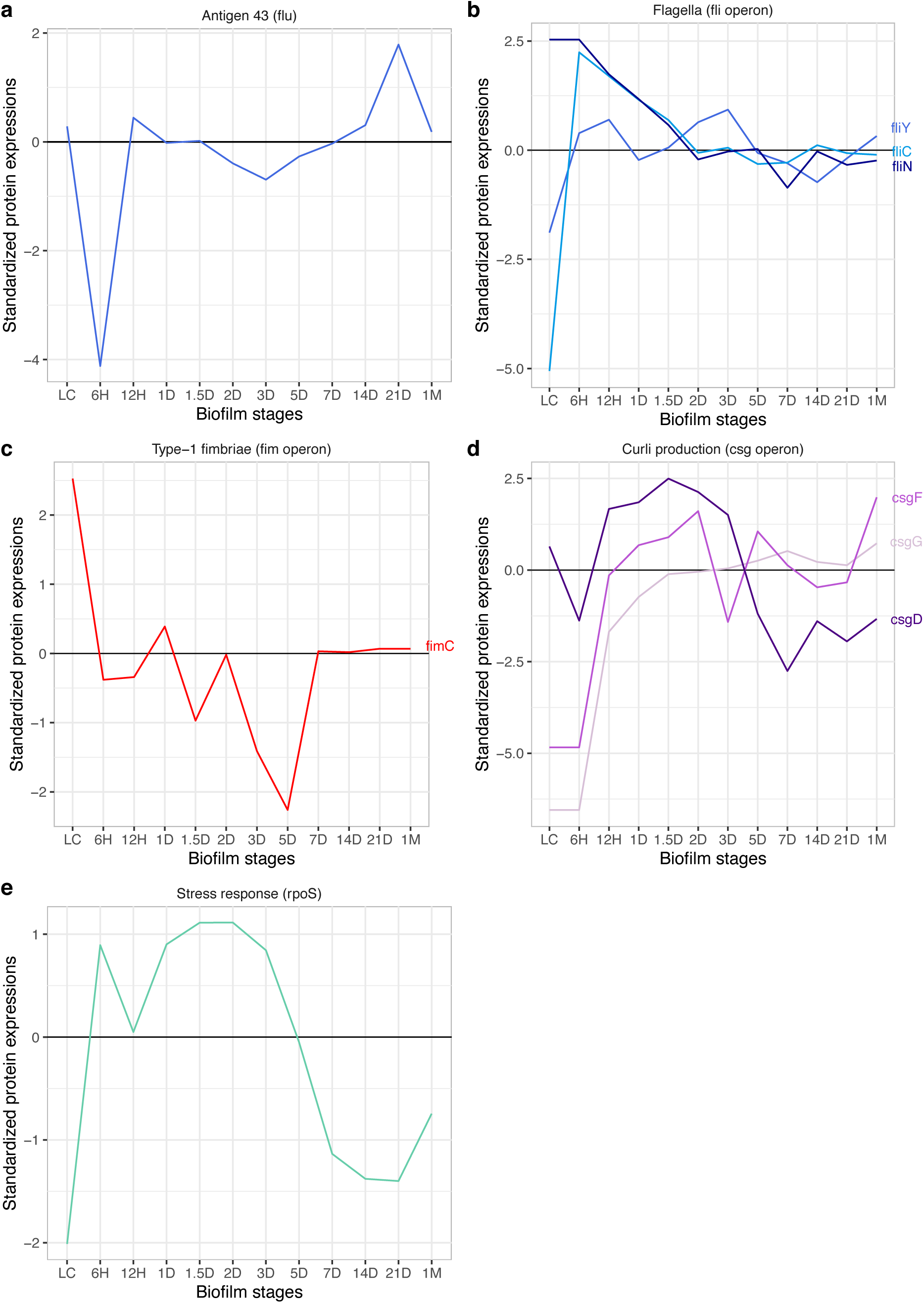
Standardized protein profiles of key proteins involved in rosette-initiated biofilm development: (**a**) Antigen 43 (*flu*) (n = 1), (**b**) flagella (*fli* operon) (n = 3), (**c**) type-1 fimbriae (*fim* operon) (n = 1), (**d**) curli production (*csg* operon) (n = 3), and (**e**) stress response (*rpoS*) (n = 1). Proteins encoded by *ycdSRQP* operon (polysaccharide matrix) were not detected in our proteome data. Grey horizontal line represents the median of standardized proteome expression values. For a full list of proteins used in profile visualization and their corresponding expression values, refer to Supplementary File S14.

